# Next generation statistical framework for next generation spatial transcriptomics data

**DOI:** 10.1101/2025.05.02.651852

**Authors:** Fatoumata Mangane, Pierre Bost

## Abstract

The rapid advancement of spatial transcriptomic technologies, particularly in situ hybridization based methods, has enabled the profiling of gene expression at sub cellular resolution across large tissue sections. Commercial platforms such as Xenium and CosMx now routinely generate high-quality datasets of increasing size and complexity. However, existing analytical approaches, often adapted from single-cell genomics, fall short in addressing the specific challenges posed by spatial data, especially at scale. In this work, we present TranspaceR, a new R package that introduces computational and statistical methods tailored to the analysis of next-generation spatial transcriptomic datasets. Our framework includes novel quality control procedures, scalable gene selection strategies especially for spatially variable genes, and optimized normalization and dimensionality reduction techniques based on in-depth statistical characterization of spatial data. We also demonstrate how single-cell annotation tools can be leveraged for automated cell-type labeling within spatial contexts. Together, these tools enable the efficient and robust analysis of imaging-based spatial transcriptomics datasets comprising millions of cells, paving the way for deeper insights into tissue organization.

## Introduction

In recent years there has been a meteoric development of spatial transcriptomic technologies. Among them, in-situ hybridization based methods can profile the expression of hundreds of genes with a sub-cellular resolution, allowing the characterization of the spatial structures of health and diseased tissues with unprecedented scale [Moffit et al.]. The recent release of commercial platforms such as Xenium, CosMx or MERSCOPE now enables the generation of high-quality datasets for which the number and size are increasing constantly and exponentially [Moses and Pachter].

However, despite these significant technological improvements, the analysis of such datasets remains a notable challenge and optimal strategies for quality control, data normalisation, and cell clustering have yet to be identified as so far those are derived from the single-cell genomic fields [Atta et al.] and thus do not account for the specificities of spatial transcriptomics data. In addition, most of the methods developed for dedicated spatial analysis, such as the detection of spatially variable genes, have been developed for small-scale datasets and thus cannot be applied to the million cell large datasets [Chen et al.]. It is thus necessary to develop new computational approaches and tools to efficiently analyse this new generation of spatial transcriptomic datasets which has the potential to change our understanding of tissue organization.

In this paper, we introduce tailored computational and statistical methods for the analysis of next-generation in situ hybridization-based spatial transcriptomics techniques, and present them in a new R package, TranspaceR. These methods notably includes new quality control strategies and gene selection methods, including highly scalable methods for the detection of spatially variable genes. In addition, we performed an in-depth analysis of spatial transcriptomic data statistical properties to determine the optimal normalisation and low-dimensional embedding strategies, resulting in a significantly improved analysis. Finally, we have shown that single-cell annotation tools can be used to drastically facilitate the analysis of spatial transcriptomic datasets and directly annotate individual cells.

Altogether we provide robust and scalable statistical approaches that can analyze imaging based ST datasets containing millions of cells in a few minutes, thus facilitating the analysis of next-generation ST datasets.

## Results

### Next generation spatial transcriptomic data represents a new analytical challenge

In order to better understand what are the challenges associated with the data generated by new generation spatial transcriptomic techniques such as Xenium and CosMX we analyzed the temporal evolution of studies referenced in the Museum of Spatial Transcriptomic database [Moses and Pachter] (Methods). We first observed an exponential increase in the number of published studies with more than 150 studies being published in 2023, illustrating the popularization of spatial transcriptomic techniques (Figure 1A). In addition to being more numerous, spatial transcriptomic datasets also tend to become significantly bigger over time, with recent datasets having on average more than 10^5^ cells (Figure 1B) in contrast to the first datasets that rarely described more than 1000 cells. In comparison, the number of measured genes only slightly yet significantly, increased over time (Supplementary Figure 1A). Finally, we observed a clear shift regarding the type of samples studied by spatial transcriptomics: while in 2020 the majority of samples described by spatial transcriptomics were obtained from healthy mouse samples, in 2024 half of the samples were of human origin (Supplementary Figure 1B) or were not considered ‘healthy’ (Figure 1C). This change is likely driven by the recent appearance of commercial platforms, which now represent more than a third of all datasets produced in 2024 (Supplementary Figure 1C). Altogether, our analysis of the Museum of Spatial Transcriptomic metadata reveals that the nature of spatial transcriptomic data have shifted from small datasets describing healthy mouse tissues to large datasets describing healthy and diseased human and mouse samples. Therefore, in order to analyze this new generation of complex datasets, new scalable and robust computational tools are needed. Throughout this manuscript, we will propose various approaches and will benchmark them on distinct datasets (Figure 1D and E, see Methods for more details) including a Xenium dataset derived from a healthy human lymph node, a CosMX dataset obtained from a human healthy prefrontal cortex and a MERFISH dataset generated from a human healthy tonsil [Zhao et al.].

**Figure 1:**
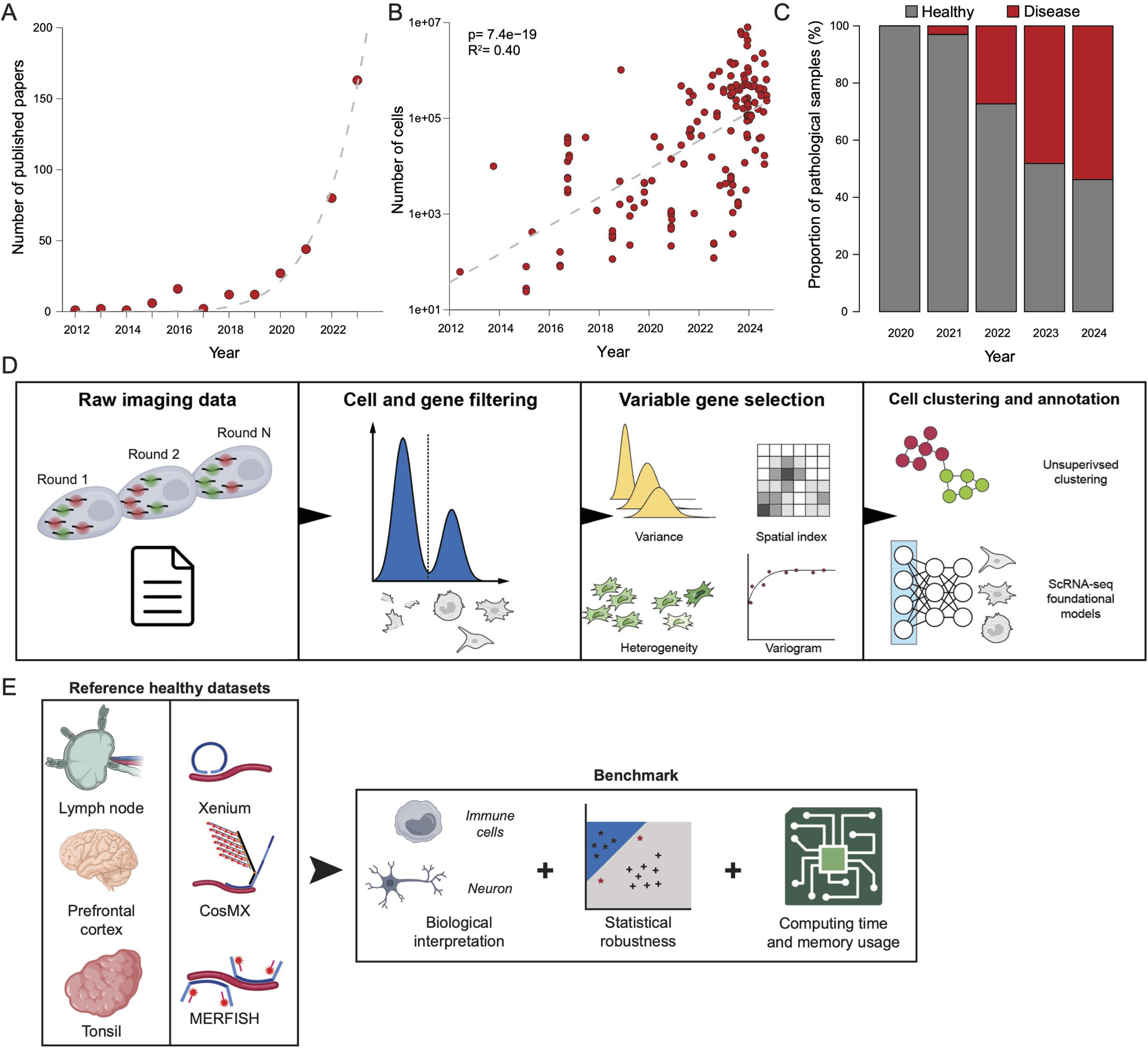
**(A)** Number of paper containing multiplexed smFISH data published each year according to the Museum of spatial transcriptomic. The grey dashed curve corresponds to an exponential fit. **(B)** Number of cells measured in multiplexed smFISH datasets according to publication year. The grey dashed line corresponds to a linear fit in the logarithmic space. Statistical significance was assessed using a Likelihood Ratio Test. **(C)** Proportion of multiplexed smFISH datasets containing non-healthy/diseased samples across time. **(D)** Workflow of the TranspaceR package. **(E)** Datasets and type of benchmarks used in the paper.

### Low quality gene and cell filtering

The filtering of low quality observations and features is the key first but also one of the most important steps of any data analysis pipeline [Hong et al.]; in the context of cell-based ST data this consists in the removal of low quality cells, generally the result of improper cell segmentation, and of lowly expressed genes. We implemented two complementary strategies for the removal of low quality cells: first, we reasoned that low quality cells would display an abnormal cell volume and therefore tried to identify a clear upper and lower bound for a correct cell radius (Figure 2A, Supplementary Figure 2B and 2C). However, an in-depth literature analysis of the cell radius of various cell types confirmed that cell size is highly variable (Supplementary Figure 2A) and spans over one order of magnitude. Therefore, the upper limit for cell size has to be set differently for each dataset, according to the tissue histology: for instance, while no cells with a radius bigger than 10µm is expected in lymphoid tissues, hepatocytes, the predominant liver cell type, have a radius that can exceed 15µm and would be filtered out with the 10µm threshold. The table we provide might thus help readers to determine the optimal threshold for their own datasets (Supplementary Table 1). Our second strategy relies on the removal of cells with a low number of detected RNA molecules, similarly to what is done for single-cell RNAseq (scRNA-seq) data: we considered that the commonly-used threshold in the field of scRNA-seq of 1000 unique RNA molecules detected could be adapted to ST datasets by scaling the threshold based on the gene panel size (Methods). Through the manuscript we have systematically used both strategies to be sure that no low-quality cells are kept for further analysis.

**Figure 2:**
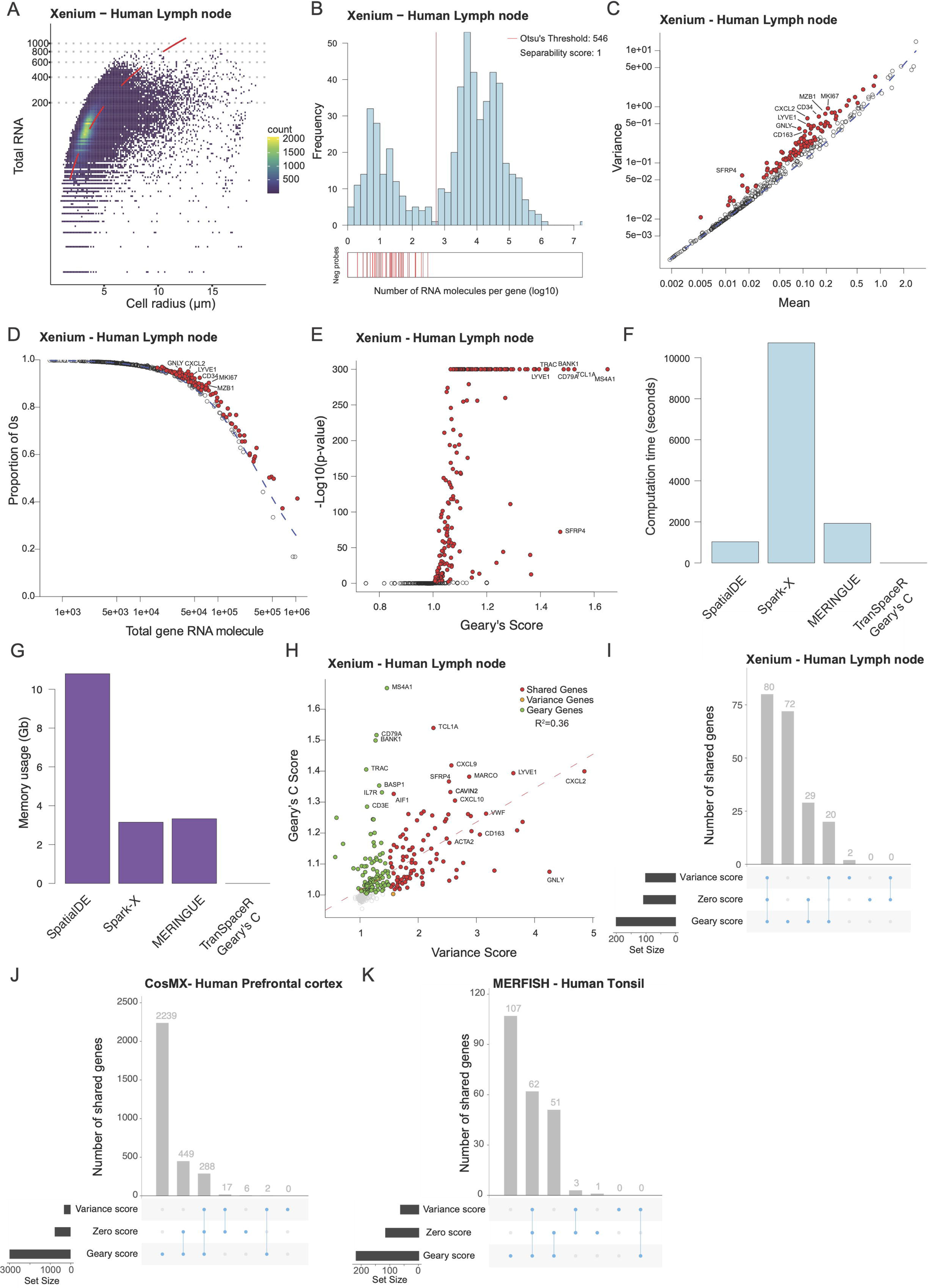
**(A)** Relationship between cell radius and total RNA molecules detected in the Xenium human lymph node dataset. The red dashed line corresponds to a quadratic fit. **(B)** Distribution of the number of RNA molecules detected for each gene in the Xenium human lymph node dataset. The vertical red line corresponds to the threshold computed using Otsu’s method. **(C)** Relationship between gene expression mean and variance in the Xenium human lymph node dataset. The dashed blue line corresponds to the theoretical relation between gene expression mean and variance according to the fitted negative binomial model. Genes displaying a significant excess of variance (corrected p-value <0.01) are colored in red. **(D)** Relationship between gene expression mean and proportion of zeros in the Xenium human lymph node dataset. The dashed blue line corresponds to the theoretical relation between gene expression mean and proportion of zeros according to the fitted negative binomial model. Genes displaying a significant excess of zeros (corrected p-value <0.01) are colored in red. **(E)** Geary’s C score for gene expression in the Xenium human lymph node dataset. Genes displaying a significantly high Geary’s C score (corrected p-value <0.01) are colored in red. **(F)** Computation time needed to analyze a subset of 10.000 cells from Xenium human lymph node dataset for various algorithms. **(G)** Memory usage when analyzing a subset of 10.000 cells from Xenium human lymph node dataset using various algorithms. **(H)** Comparison of the variance and Geary’s C score for the Xenium human lymph node dataset. The dashed red line corresponds to a linear fit. **(I)** UpSet plot comparing variable genes identified using the variance, zero and Geary’s C score for the Xenium human lymph node dataset. **(J)** UpSet plot comparing variable genes identified using the variance, zero and Geary’s C score for the CosMX human prefrontal cortex dataset. **(K)** UpSet plot comparing variable genes identified using the variance, zero and Geary’s C score for the MERFISH human tonsil dataset.

Regarding lowly expressed genes, their removal was performed using two complementary methods: automated thresholding, and background estimation by negative probes (Figure 2B). The first method consists in computing a general expression threshold by Otsu’s method [Otsu 1979] to separate unexpressed or lowly expressed genes from expressed genes (Methods). In opposition, the second method relies on the use of ‘negative probes’ (probes that should not be detected as they typically target genes from an other species) to define an expression threshold (Figure 2B, Figure S2D and S2E, Methods). We realized that the quality of datasets can be assessed by looking at how both methods agree by computing the proportion of negative probes which are considered as being ‘un-expressed’ according to Otsu’s method, and therefore a score closer to 1 corresponds to a high quality dataset (Figure 2B, Figure S2D and S3E).

### Parametric and statistically robust gene selection

While single-cell and spatial transcriptomic can now profile more than thousands of genes, only a part of them are usually kept for analysis as a significant fraction of genes do not display biologically relevant variations, such as cell-type or condition-specific expression. Therefore, selecting truly biologically variable genes is an essential step of spatial transcriptomic analysis [Yang et al.].

It is worth mentioning that n contrast to scRNA-seq data, only a limited number of genes are being measured in cell-based ST experiments. There is thus a need for a robust gene selection method based on a statistical model where the number of variable genes is not predefined. Accordingly, we make the hypothesis that a non-variable gene expression follows a negative binomial distribution which over-dispersion parameter is shared across all genes. First we compute the shared over-dispersion parameter by fitting the global variance-mean relationship using a simple linear model. Using this parameter, we compute the theoretical variance for each gene and compared it to the observed variance before computing a corresponding p-value through asymptotic estimation (Figure 2C, Figure S2F and S2G, Methods). We implemented a similar approach that models the proportion of zeros (sometimes improperly called ‘drop-out’ in the single-cell field) to identify genes displaying an abnormally high proportion of zeros (Figure 2D, Figure S2H and S2I Methods). When applied to the Xenium human lymph node dataset (Figure 2E), both methods correctly identify biologically relevant genes as being the most variable, including markers for cell division (MKI67), cytotoxic T cells (GNLY), plasma cells (MZB1) and lymphatic endothelial cells (LYVE1) markers. Similarly, when applied to the CosMX human prefrontal cortex (Supplementary Figure 2J), our methods identified key marker genes of neurons (NPY and NRGN), astrocytes (SLC1A2, AQP1 and AQP4) and oligodendrocytes (PLP1) populations, as well as when applied to the MERFISH human tonsil dataset with the detection of B-cells (TCL1A), neutrophils (MMP9), cytotoxic T-cells (CD8A), NK cells (NKG7) and plasma cells (IGHD) markers (Supplementary Figure 2K).

Complementary to these two non-spatial approaches, we developed a highly efficient implementation of the Geary’s C spatial index [Cliff and Ord], a commonly used but poorly scalable measure of spatial autocorrelation, by reformulating the equation and the combined use of optimized Delaunay triangulation implementation and sparse matrix operation (Figure 2E, Figure S2J and S2K, Methods). We then decided to compare our Geary’s C implementation to three commonly used spatially variable gene detection methods: SpatialDE [Svensson et al.], a Gaussian process based approach, Spark-X [Zhu et al.] a non-parametric method, and MERINGUE [Miller et al. 2021], a Geary’s C index based method. We ran all four methods on a 10.000 cell subset of the Xenium lymph node dataset and observed that while our Geary’s C index implementation requires less than one second and less than 1Mb of memory (Figure 2F and G), other methods required between 15 minutes to 3 hours and 3 to 11Gb of RAM to run, making them incompatible with large scale spatial transcriptomic datasets.

Interestingly, the top genes identified by the Geary’s C score differ from the ones identified by both the variance and zero score as they are mostly B cell markers such as MS4A1 (encoding the CD20 gene), TCL1A, CD79A and BANK1; moreover only a limited correlation was observed between the variance and Geary’s C score (Figure 2H). We thus investigated the overlap between the genes identified by each method for the lymph node dataset and noted that while the majority of gene spicked up by the variance and zero method were also detected according to Geary’s C score, a significant number of genes were specifically detected by the last method (Figure 2I). We made a similar observation when we applied the three methods to a human prefrontal cortex dataset generated by the CosMX technology (Figure 2J). However, all three methods provided highly concordant results for a mouse liver dataset produced using the MERFISH technology (Figure 2K), suggesting that depending on the dataset the agreement between methods can widely vary. As a general strategy, we consider that the union of all three methods results will provide a consistent and biologically meaningful gene list for further analysis.

In sum, we introduced three robust and statistically founded methods for the detection of variable genes, including a novel highly scalable method for the detection of spatially variable genes.

### Fast Fourier Transform for scalable variogram computation

While Geary’s C is an informative measurement it suffers from two limitations: first it is highly dependent on the definition of spatial proximity, and secondly it does not provide any quantitive information on the studied spatial structure such as the scale of the spatial autocorrelation. A possible alternative to Geary’s C and other similar indexes is the empirical variogram, a mathematical function computed from the data that describes the average difference of measured values located at a certain distance [Cliff and Ord]. At a given distance, a low variogram value corresponds to a strong spatial autocorrelation while a higher value is suggesting a low spatial autocorrelation. Following their computation, empirical variograms are usually interpreted by fitting various variogram models and identifying the best theoretical variogram model. While being highly informative this approach is rarely used in practice due to the prohibitive computational cost required to compute pairwise differences for each pair of point for large scale datasets.

In order to overcome this challenge and make the computation of empirical variograms possible for large-scale spatial transcriptomic dataset, we introduce a new strategy based on rasterization and matrix convolution. Indeed, when the data points are localized on a regular grid variogram computation can be seen as a convolution product and therefore can be computed by fast Fourrier transform (FFT) [Marcotte]. Our approach thus consists in the ‘rasterization’ of the data, i.e. the transformation of irregular spatial data into a matrix by pooling the expression of neighboring cells, before computing the empirical variogram by FFT, then finally identifying the theoretical variogram that fits best to the empirical variogram (Figure 3A) (Methods). More precisely we use three variogram models: a constant model (corresponding to an absence of spatial autocorrelation), an exponential one and what we called a fine-tuned exponential model (Methods).

**Figure 3:**
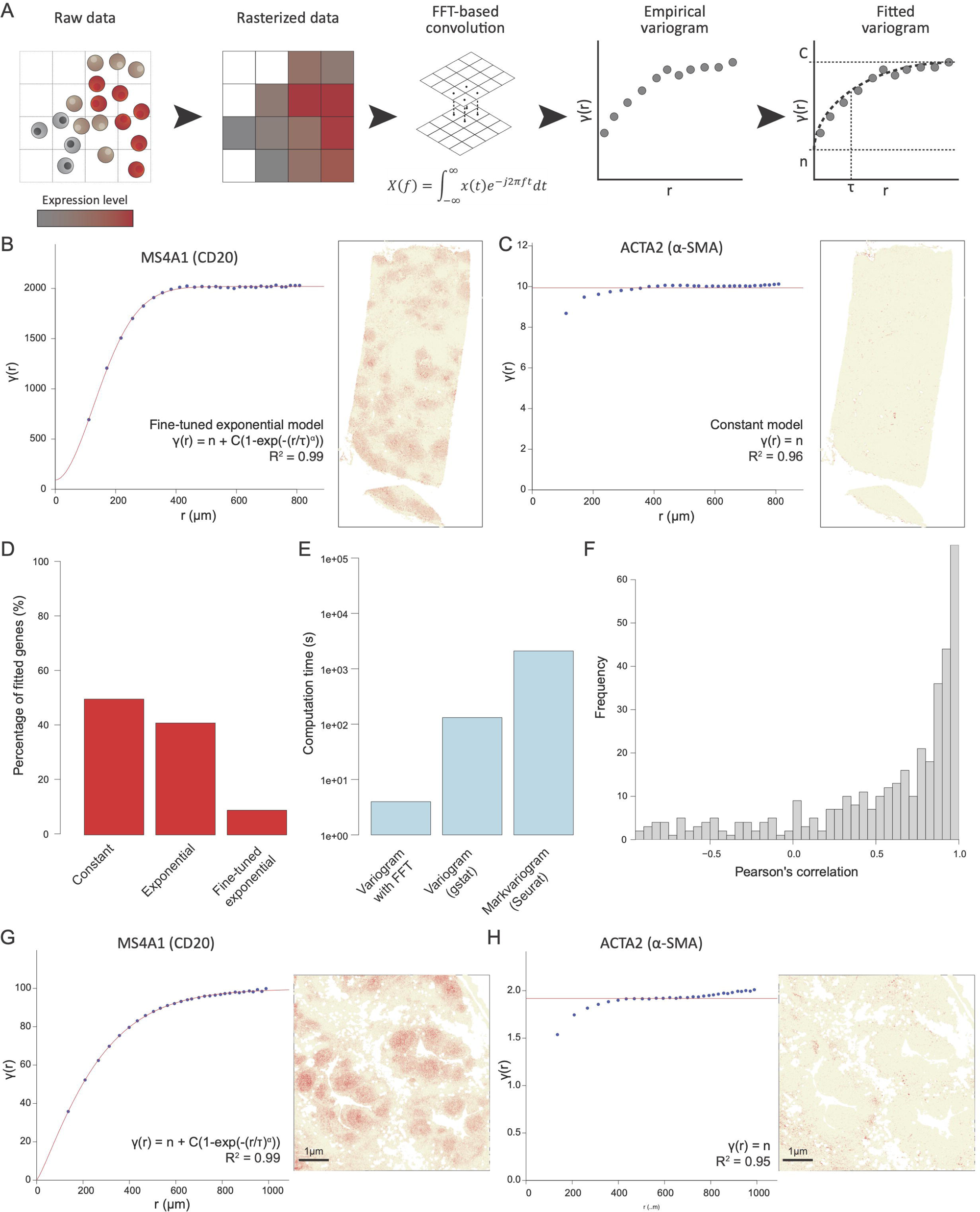
**(A)** Workflow of the FFT Variogram approach. **(B)** Variogram of the MS4A1 gene encoding the CD20 marker in the Xenium human lymph node dataset. The red curve corresponds to the fitted theoretical variogram. **(C)** Variogram of the ACTA2 gene encoding the alpha smooth muscle actin protein in the Xenium human lymph node dataset. **(D)** Results of variogram fitting on the Xenium human lymph node. The red curve corresponds to the fitted theoretical variogram. **(E)** Computation time of three different variogram implementations applied to a subset of 10.000 cells of the Xenium human lymph node dataset. **(F)** Distribution of the correlation between gene expression classical variogram and FFT variogram. **(G)** Variogram of the MS4A1 gene encoding the CD20 marker in the Visium HD human lymph node dataset. The red curve corresponds to the fitted theoretical variogram. **(H)** Variogram of the ACTA2 gene encoding the alpha smooth muscle actin protein in the Visium HD human lymph node dataset. The red curve corresponds to the fitted theoretical variogram.

For instance, when applied to the B-cell marker CD20, encoded by the MS4A1 gene, in the human lymph node sample, the variogram revealed a strong spatial autocorrelation for distance values below 400µm (Figure 3B left panel) in agreement with the observed spatial pattern (Figure 3B right panel). In opposition, the variogram of the alpha smooth muscle actin gene, encoded by the ACTA2 gene, revealed limited spatial autocorrelation (Figure 3C left panel), in agreement with the biology of the gene which is supposed to be specifically expressed around blood vessels (Figure 3C right panel). Overall, out of the 436 genes studied, 50% were considered to have a flat variogram and thus not displaying any significant spatial variation while 41% and 9% of the genes were determined to have an exponential and fine-tuned exponential, respectively (Figure 3D).

Interestingly, we observed that while the genes classified as displaying a flat or exponential variograms do not display any significant enrichment of cell type markers (Figure S3A and B), genes with a fine-tuned variogram were significantly enriched in markers of various immune cells including B-cells, T-cells, plasma cells, plasmacytoid dendritic cells and regular dendritic cells (Figure S3C). Thus, our variogram approach is able to identify key biologically-relevant genes in an unsupervised manner. We then compared our FFT-based implementation to two regular and commonly used variogram computation implementations in R: one from the gstat [Pebesma et al.] package and one from the Seurat package [Butler et al.]. While for 10.000 cells and 436 genes only 4 seconds are needed with our FFT-based implementation, more than 2 and 35 minutes are required for the gstat and Seurat implementation respectively (Figure 3E). The observed trend became even more apparent when applying the methods on 300.000 cells: while only 13 seconds were required with our implementation, more than 8 hours were required with the gstat implementation, highlighting the usefulness of our approach. We then investigated how the variogram estimated by FFT was similar to the one computed by the gstat package: we observed an overall very strong correlation with a median correlation of 0.74 (Figure 3F), validating the validity of our approach.

Finally we checked whether our approach could be applied to types of spatial data other than cell-based ST. We first focused on a human tonsil dataset generated using the recently released Visium HD platform, a spot-based spatial transcriptomic technique with a cellular resolution (4µm). Similarly to the Xenium data, our FFT-variogram was able to detect highly spatially variable genes such as CD20 (Figure 3G) while a gene displaying spatial autocorrelation at a low-scale such as the alpha smooth muscle actin is considered to have a constant variogram (Figure 3H). Interestingly we observed a similar fitted variogram model for the CD20 gene across the two datasets with a range value of 153 and 197µm respectively, suggesting that our FFT-variogram is a robust measure of autocorrelation scales.

### Volume normalisation generates significant statistical artefacts

Data normalization has been a long-standing challenge in the field of scRNA-seq data [Vallejos et al.], and still represents a major issue for spatial transcriptomics. Very elegant work from the Raj team have shown that biologically, cell identity is not determined by the absolute number of mRNA molecules but rather by their concentration [Padovan-Merhar et al.]. In addition, recent works have shown that normalizing cell-based ST data by the total RNA amount can lead to significant and problematic biases when the observed gene panel is not representative [Atta et al.]. Therefore, it is tempting to directly normalize RNA expression by the cell volume for spatial transcriptomic data. Here we assume that the cell area measured by the cell segmentation step is an acceptable proxy to the actual cell volume. To test the validity of this approach we first looked at the correlation between individual gene expression and cell size across four datasets generated by the Xenium and CosMX technologies encompassing different gene panel sizes and three different tissues (Figure 4A). To our surprise we only observed a very limited correlation between the cell volume and gene expression across the four datasets (Figure 4A), suggesting that cell volume is not the major source of gene expression variation. In order to understand this phenomenon we focused on the whole-transcriptome CosMX Pancreas dataset in order to avoid biasing our analysis due to a skewed gene panel choice. Curiously, we observed a strong correlation between the expression/ volume correlation and the mean expression of the genes (Figure 4B, R2=0.79) suggesting that the expression/volume correlation can be mostly explained by the gene mean expression value.

**Figure 4:**
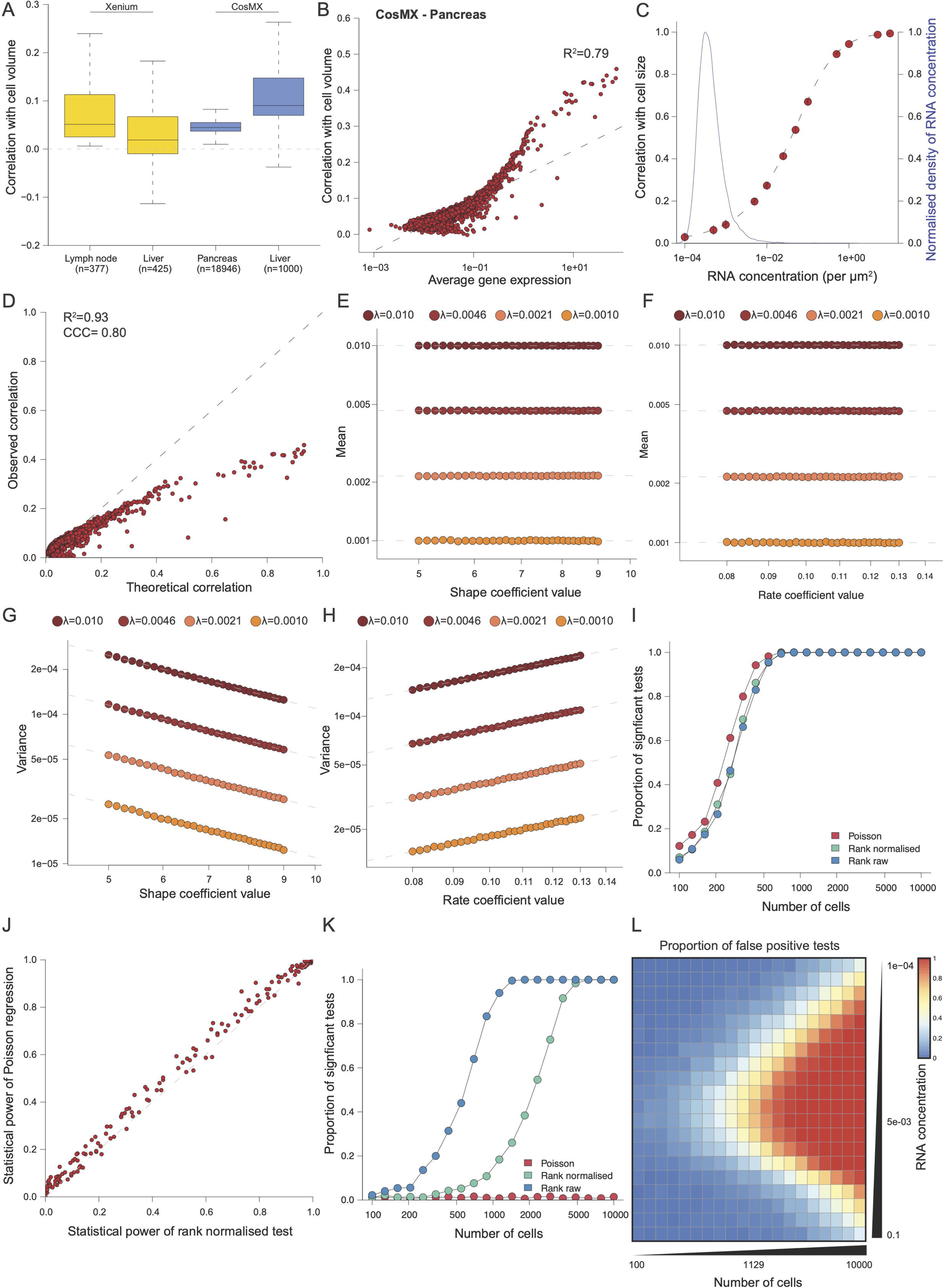
**(A)** Distribution of Pearson’s correlation between gene expression and cell volume in four district datasets. The thick line corresponds to the median, and the bottom and upper limits of the box correspond to the first and third quartiles, respectively. The lower and upper whiskers correspond to the lowest and highest values, respectively, within the range of the first and third quartiles ±1.5 times the interquartile range (IQR). **(B)** Relationship between average gene expression and correlation between gene expression and cell volume for the CosMX human pancreas dataset. The dashed grey line corresponds to a linear regression. **(C)** Correlation between cell size and gene expression as a function of RNA concentration obtained by simulation (red points) or according to the Poisson-Gamma model (dashed black line). The blue plain curve corresponds to the RNA concentration distribution observed in the CosMX human pancreas dataset. Black vertical bars correspond the standard deviation across simulations. **(D)** Comparison of the observed and theoretical/expected correlation between gene expression and cell volume. The dashed grey line corresponds to the line of equation x=y. **(E)** Relationship between the mean of synthetic volume-normalized data as a function of the shape coefficient value. Each dashed grey line corresponds a linear regression for a different RNA concentration. **(F)** Relationship between the mean of synthetic volume-normalized data as a function of the rate coefficient value. Each dashed grey line corresponds a linear regression for a different RNA concentration. **(G)** Relationship between the variance of synthetic volume-normalized data as a function of the scale coefficient value. Each dashed grey line corresponds a linear regression for a different RNA concentration. **(H)** Relationship between the variance of synthetic volume-normalized data as a function of the rate coefficient value. Each dashed grey line corresponds a linear regression for a different RNA concentration. **(I)** Power analysis of Poisson regression and of rank test on raw and volume normalized data. **(J)** Comparison of statistical power for a rank test on volume normalized data and Poisson GLM. **(K)** Specificity analysis of Poisson regression and of rank test on raw and volume normalized data. RNA concentration is fixed in this analysis. **(L)** Impact of cell number and RNA concentration on the proportion of false positive tests.

In order to investigate the causes of this relationship, we simulated gene expression values as being the product of two successive random processes: random cell size and Poisson sampling noise (see Methods for more details). Similarly to real data, we observed a monotonic sigmoid relationship between the mRNA concentration of a gene (i.e. its normalized mean gene expression) and the expression/volume correlation (Figure 4C, red points). In addition, according to this simulation, the range of observed mRNA concentrations observed in our datasets should result in expression/volume correlations lower than 0.2 (Figure 4C, dashed line), similar to what is observed with the data. We therefore decided to perform a theoretical analysis of our model by assuming that cell volume followed a gamma distribution (Supplementary Figure 4A and B). We obtained the following formula linking the mRNA concentration Ⲗ and the shape parameter θ of the cell volume gamma distribution to the expression/volume correlations (see Supplementary Note 1):

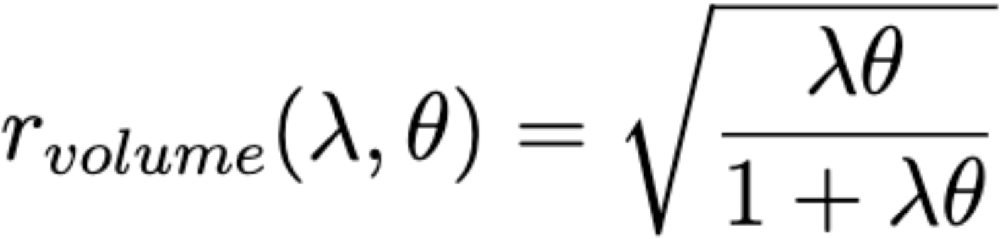

This equation shows that the correlation (and thus the percentage of variance explained) between cell volume and gene expression is the result of two opposing behaviors. For highly expressed genes (high value of Ⲗ) or highly variable volume (high θ), the main source of variation is the heterogenous cell volume while for lowly expressed genes the Poisson noise is driving inter-cell variations. This analytical expression perfectly fitted the simulated points (Figure 4C, black dashed line) and was used to check the quality of our model by predicting the expected expression/ volume correlation on the real Pancreas CosMX data. We observed a strong similarity between the observed and the predicted correlation except for highly correlating and thus highly expressed genes, thus highlighting the efficacy of our model despite its simplicity (Figure 4D, R^2^=0.93 and CCC=0.80).

As we have shown that volume variation only explains a minor fraction of the data variation, we wondered what would be the effect of normalizing the data by dividing gene expression values by the cell volume. To achieve this, we generated synthetic data by first sampling a cell volume from a gamma distribution with realistic parameters, then applying Poisson sampling, and finally normalizing the data by dividing it by the cell volume (Supplementary Figure 4C). We generated data with different parameters for the cell volume distribution and looked at the impact on the mean and variance of the generated data: while the mean did not seem affected by changes of the shape and rate parameters (Figure 4E and F), the variance was strongly impacted by both parameters (Figure 4G and 4H). Similarly, higher moments such as the skewness and the kurtosis were also affected (Supplementary Figure 4D to G) validating the fact that while the volume normalisation is efficient at correcting the mean, it is unable to correct volume-bias for higher moments.

Finally, we wondered what would be the impact of volume normalisation on differential gene expression analysis. To do so, we first simulated gene expression values of two different cell populations, one having an RNA concentration set to be twice that of the other cell population while the cell volume distribution remains the same for both populations. The simulated values were used to perform a power analysis using either a rank test on the raw values, a rank test on the volume normalized values, or a Poisson Generalized Linear Model (GLM) with the cell volume as a covariate (Methods). We first compared the statistical power of all three methods for a given RNA concentration value (λ=0.01) and observed that all three test performed similarly, with the Poisson GLM performing only slightly better (Figure 4I) compared to both rank tests. We then performed a more systematic analysis by varying the reference RNA concentration level and observed that on average, the Poisson test indeed displayed a moderate albeit significant gain of statistical power over a rank test on the volume normalized values (Figure 4J, Supplementary Figure 4H and I, paired t-test: p=4.9e-22). We then wondered whether these tests differed in their robustness and thus simulated gene expression of two identical cell populations (identical volume distribution and RNA concentration) before performing differential expression analysis from which we observed a similar and well-controlled False Positive Rate (FPR) for all three methods (Supplementary Figure 4J) . However, to our surprise, when the cell volume distribution parameters were slightly changed (25% increase of the shape parameter value) we observed a FPR that increases with the number of sampled cells for the rank test on raw data, but also for the rank test performed on the volume normalized data (Figure 4K and L). In opposition, the Poisson GLM test was robust to changes of the cell volume distribution, suggesting it is able to properly remove the effects of volume variation unlike the rank tests. Those tests are indeed known to be sensitive to changes in distribution moments other than the mean, especially when the number of samples is reaching a certain limit [Fagerland].

Altogether our analysis revealed that volume is not the main source of non-biological noise in spatial transcriptomic data and that normalizing spatial transcriptomic data by the volume is not adequate and can result in significant statistical artifacts.

### Optimal low dimensional embedding for unsupervised clustering

Low dimensional embedding is an essential step in any single-cell analysis as it allows for the aggregation of correlated gene expression measurement in order to mitigate the intrinsic noise of single-cell measurements and thereby improve further analysis steps such as unsupervised clustering (Figure 5A) [Xiang et al.]. Additionally, in the context of targeted ST, we hypothesized that some gene panels might be highly unbalanced with a few cell types benefiting from the majority of the markers within the panel, resulting in a strongly biased analysis toward these cell types, and therefore dimensionality reduction could also reduce this bias.

**Figure 5:**
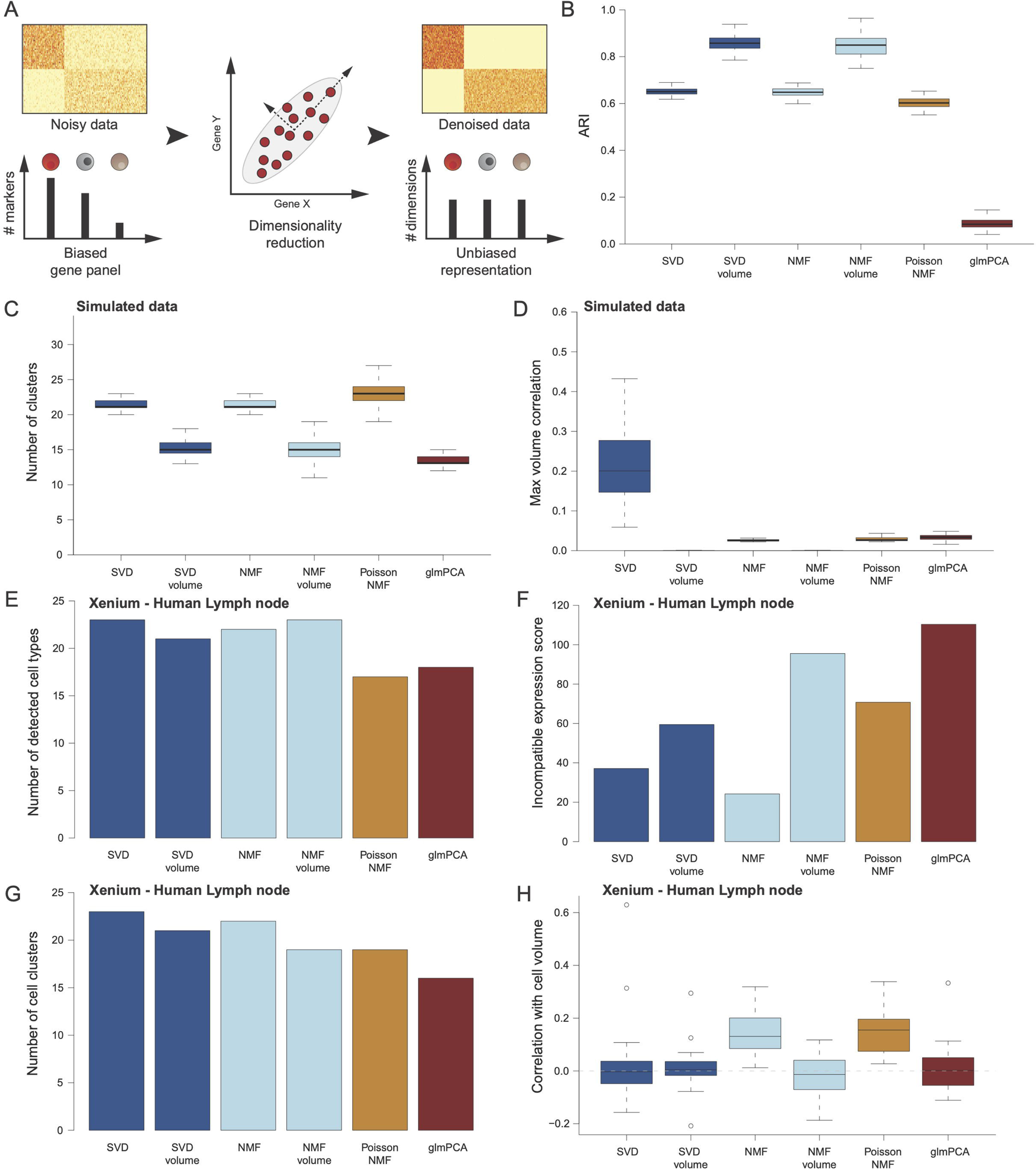
**(A)** Motivation for the use of dimensionality reduction techniques. **(B)** Distribution of the Adjusted Rand Index (ARI) between inferred and reference clusters according to the dimensionality reduction technique used when applied to simulated data. The thick line corresponds to the median, and the bottom and upper limits of the box correspond to the first and third quartiles, respectively. The lower and upper whiskers correspond to the lowest and highest values, respectively, within the range of the first and third quartiles ±1.5 times the IQR. The boxplot is based on N=300 simulations. **(C)** Distribution of the number of clusters inferred according to the dimensionality reduction technique used when applied to simulated data. The thick line corresponds to the median, and the bottom and upper limits of the box correspond to the first and third quartiles, respectively. The lower and upper whiskers correspond to the lowest and highest values, respectively, within the range of the first and third quartiles ±1.5 times the IQR. The boxplot is based on N=300 simulations. **(D)** Distribution of the maximal correlation between inferred latent dimensions and cell volume according to the dimensionality reduction technique used when applied to the Xenium lymph node dataset. The thick line corresponds to the median, and the bottom and upper limits of the box correspond to the first and third quartiles, respectively. The lower and upper whiskers correspond to the lowest and highest values, respectively, within the range of the first and third quartiles ±1.5 times the IQR. The boxplot is based on N=300 simulations. **(E)** Number of cell types detected according to enrichment analysis according to the dimensionality reduction technique used when applied to the Xenium lymph node dataset. **(F)** Average incompatible expression score according to the dimensionality reduction technique used when applied to the Xenium lymph node dataset. **(G)** Number of cell clusters detected according to the dimensionality reduction technique used when applied to the Xenium lymph node dataset. **(H)** Distribution of the correlation between inferred latent dimensions and cell volume according to the dimensionality reduction technique used when applied to the Xenium lymph node dataset. The thick line corresponds to the median, and the bottom and upper limits of the box correspond to the first and third quartiles, respectively. The lower and upper whiskers correspond to the lowest and highest values, respectively, within the range of the first and third quartiles ±1.5 times the IQR. The boxplot is based on N=20 latent dimensions.

In order to determine the optimal dimensionality reduction techniques for ST data, we first benchmarked a list of methods on simulated ST data: using the statistical framework described in the previous section we simulated 10 cell clusters with a variable number of marker genes of variable expression level (see Methods for details). We then performed dimensionality reduction before performing Louvain’s graph based clustering [Levine et al. 2015] and compared the obtained clustering with the initial cluster labels. We compared 6 methods that are commonly used for scRNA-seq data analysis : Singular Value Decomposition (SVD) on un-normalized and volume normalized data, Non negative Matrix Factorisation (NMF) on un-normalized and volume normalized data [Debruine et al.], Poisson NMF [Lee and Seung], and Generalized Principal Component Analysis with a Poisson posterior distribution (glmPCA) [Weine et al., Townes et al.] (Methods). To our surprise we observed that the SVD and NMF performed on the volume normalized data exhibited the best results with the highest Normalized Mutual Information (NMI) score and Adjusted Rand Index (ARI) (Figure 5B, Figure S5A). Similarly, while the number of clusters was over-estimated for all methods, a common problem with Louvain’s clustering, both SVD and NMF performed on normalized data displayed the lowest number (Figure 5C). We then wondered whether some of the dimensionality reductions were driven by cell volume variation and observed that while SVD performed on the raw data displayed the highest maximal correlation the maximal correlation did not exceed 0.2 on average (Figure 5D). In contrast, all other methods displayed a maximal correlation with cell volume that was below 0.05.

We then tested the six approaches on the human Xenium lymph node dataset. As we do not have a reference cell annotation we could not compute an ARI or NMI score and decided instead to check whether the identified clusters were biologically relevant. Using the PanglaoDB marker database [Franzén et al.] to perform Gene Set Enrichment Analysis (GSEA) (Methods) we computed the number of different cell types that were detected for each method: we observed a similar number of detected cell types for the SVD and NMF based clustering (between 21 to 23 cell types, Figure 5E) while the Poisson NMF and the glmPCA derived clustering only resulted in 17 and 18 detected cell types. We then wondered whether some approaches would tend to group biologically dissimilar cells in the same cell cluster. We thus manually curated the dataset gene panel and annotated genes into 11 mutually exclusive categories: B-cells (BANK1, CD19, MS4A1, TNFRSF13B and TNFRSF17), Dendritic Cells (LAMP3, CCR7), blood endothelial cells (PECAM1 and VWF), lymphatic endothelial cells (LYVE1, MARCO and PROX1), monocytes/macrophages (CD14, CD163, CD68, MS4A6A and MRC1), mast cells (CPA3, CTSG, MS4A2 and SLC18A2), NK cells (KLRB1, KLRC1, KLRD1 and NKG7), plasmacytoid Dendritic Cell (IL3RA, IRF8, LILRA4, PLD4 and TCF4), plasma cells (MZB1) and T-cells (CD28, CD3D, CD3E, CD4, CD8A, CTLA4, FOXP3, IL2RA, IL7R, LAG3, SLAMF1 and TRAC). Using this annotation we derived an incompatible expression score that quantifies the amount of biologically impossible co-expression patterns (Methods). To our surprise we observed that the SVD and NMF performed on raw data resulted in the lowest incompatible expression score and that the volume normalisation resulted in an increased score (Figure 5F). Moreover, both Poisson NMF and glmPCA resulted in a higher score (Figure 5F). These results could be explained by the higher number of cell clusters obtained by the ‘raw’ SVD and NMF approaches compared to the Poisson NMF and glmPCA, but also by the SVD and NMF performed on volume normalized data (Figure 5G).

Finally we wondered if some of the computed latent dimensions correspond to the cell volume and could thus confound the analysis. We thus computed the correlation between each latent dimension and the cell volume for each method and observed striking differences between each approach: while latent dimensions resulting from NMF performed on raw data and Poisson NMF display an homogeneous positive correlation across dimensions (median correlation of 0.13 and 0.15 respectively, Figure 5H), the first SVD dimension strongly correlated with cell volume (R=0.63) despite the majority of dimensions not correlating with cell volume (median correlation of -0.003).

Altogether our results highlight the necessity of performing the dimensionality reduction step on raw data using either SVD or NMF in order to obtain the most biologically meaningful and exhaustive cell clustering.

### Single-cell atlases can be used to annotate next generation spatial transcriptomic data

The constant increase of available scRNA-seq datasets has enabled the development of single-cell foundational models that can be used to annotate large-scale single-cell datasets [Szałata et al.]. As such tools are powerful facilitators of single-cell data analysis we wondered whether they could be also used to analyze and annotate spatial transcriptomic datasets in a quick and comparable fashion.

We decided to test the recently published SCimilarity tool as its has been purposely designed to be robust concerning the absence of measurements of some genes [Heimberg et al. 2024]. In order to assess whether such elaborate computational methods were necessary for the automated annotation of spatial transcriptomic data, we also tested the CellTypist method [Dominguez Conde et al. 2022] that is based on L2 penalized logistic classifiers trained from references scRNA-seq datasets. We first applied both methods to the Xenium human lymph node sample, which resulted in the expected identification of immune populations as the major cell types according to both methods (Figure 6A). In parallel we compared the computational efficiency of both methods: we observed that while for cell numbers below 4.10^5, both methods displayed similar computational time, Scimilarity was significantly faster than CellTypist in the context of large datasets (Figure 6B).

**Figure 6:**
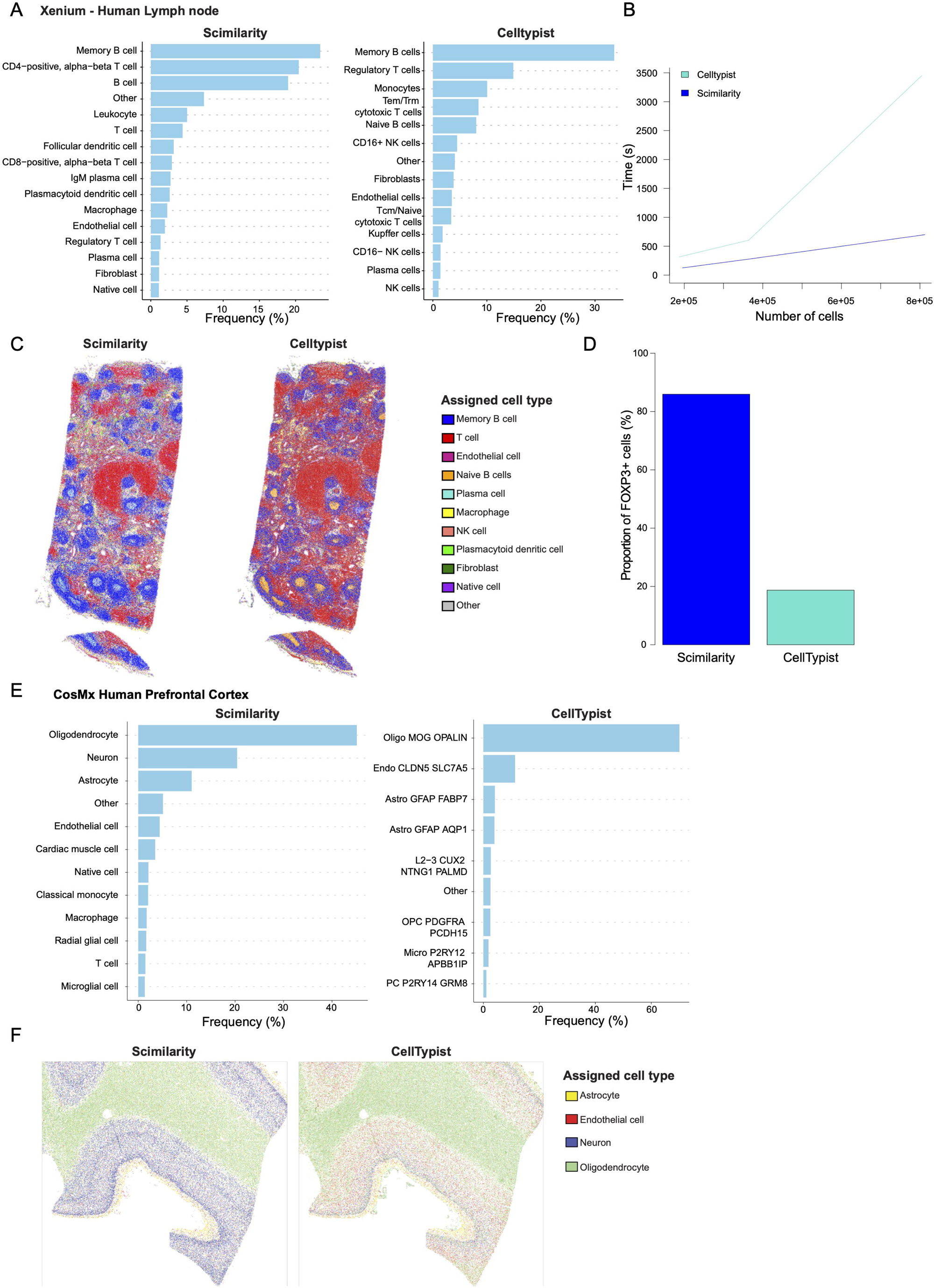
**(A)** Results of the automated annotation of the Xenium human lymph node dataset by Scimilarity (left) and CellTypist (right). **(B)** Comparison of computation time for Scimilarity and CellTypist. **(C)** Spatial structure of the Xenium human lymph node dataset according to Scimilarity (left) and CellTypist (right). **(D)** Proportion of regulatory T cells expressing FOXP3 according to Scimilarity and CellTypist. **(E)** Results of the automated annotation of the CosMX human prefrontal cortex dataset by Scimilarity (left) and CellTypist (right). **(F)** Spatial structure of the CosMX human prefrontal cortex dataset according to Scimilarity (left) and CellTypist (right).

We then decided to inspect in detail the results of both annotation approaches. While the overall tissue architecture obtained through both annotation methods is similar (Figure 6C), we observed significant anomalies with the obtained CellTypist annotation. Notably we observed an abnormally high proportion of regulatory T cells (Treg, 14.9%) which is incompatible with our knowledge of human lymph node cellular composition [Wei et al. 2006]. A more in-depth analysis confirmed that less than 20% of the cells annotated by CellTypist as Treg are expressing the canonical marker FOXP3, while more than 80% of regulatory cells annotated by Scimilarity express it (Figure 6D), validating the fact that CellTypist is incorrectly annotating Treg. Even more critically, we observed that cells located in the center of B-cell follicles were annotated as naive B cells by CellTypist, in disagreement with the known histology of these follicles that should contain activated B cells and B-cell differentiating into plasma cells (Figure 6C). In contrast, Scimilarity annotates these cells as plasma cells, which is much more coherent with the known follicular biology. To our surprise, when we compared both supervised annotations with the result of unsupervised graph clustering (Supplementary Figure 6A and B), we observed similarly low agreements between both supervised annotations and the unsupervised clustering with an ARI score of 0.27 and 0.25 for Scimilarity and CellTypist respectively.

Finally, we decided to apply the same strategy to the CosMX human prefrontal cortex data and observed that while both methods were able to identify the expected major cell types, i.e. oligodendrocytes, neurons, astrocytes and endothelial cells, cell type proportions were not in agreement between the two methods (Figure 6E). Notably, while Scimilarity annotates 45% of the cells as oligodendrocytes, 70% were annotated as such by CellTypist, a number hither than expected. Inversely, the proportion of annotated neurons was surprisingly low in the case of CellTypist annotation with only 2.6% of cells being labelled as such (L2−3 CUX2 NTNG1 PALMD) in comparison to more than 20% of cells annotated as neurons by Scimilarity, which is much closer to the actual estimate [Hannon et al.]. A closer look at the annotated tissue architecture revealed that neuron cortex layers were correctly identified by Scimilarity but not by CellTypist (Figure 6F), and were instead annotated as oligodendrocytes and endothelial cells, suggesting that deep-learning tools trained on scRNA-seq data properly annotate complex ST datasets, unlike regular machine-learning based approaches.

Altogether, our analysis demonstrated that deep-learning based annotation tools initially designed for scRNA-seq data analysis can be used to facilitate the analysis of spatial transcriptomic data and drastically accelerate the annotation of individual cells with a high resolution.

## Discussion

Here, we present the statistical foundations for the analysis of large-scale multiplexed smFISH datasets, along with TranspaceR, a novel and dedicated R pipeline designed specifically for this purpose. We first introduce a new method to assess whether a gene exhibits a statistically significant excess of variance, alongside a highly scalable implementation of Geary’s C index, both of which greatly facilitate the critical step of gene selection. To further expand the toolkit for spatial data analysis, we developed the first highly scalable implementation of the variogram, applicable to both single-cell and spot-based spatial transcriptomics. This enables a systematic, quantitative analysis of spatial gene expression patterns. Through simulations and variance decomposition, we identified Poissonian sampling—not cell volume heterogeneity, as previously assumed—as the primary source of gene expression variance. Based on this theoretical insight we devised a new normalization and clustering strategy that enhances the recovery of biologically meaningful cell clusters from spatial transcriptomics data. Finally, we validated the performance of Scimilarity, a recently published deep learning annotation tool trained on scRNA-seq atlases, for annotating spatial transcriptomics datasets.

Together, our contributions provide a comprehensive and scalable toolkit for the analysis of next-generation spatial transcriptomics data, enabling rapid, in-depth analyses on a single laptop within minutes.

Our approach and pipeline do have limitations. First, the spatial analysis tools presented in this manuscript focus solely on the behavior of individual genes, and do not account for potential interactions between gene pairs. As a result, biologically meaningful spatial relationships, such as the interaction between CD3 and CD20, which delineates T-cell and B-cell zones within lymphoid organs, are not captured by our current analysis framework. Therefore, one of the subsequent key next milestones in spatial omics analysis will be the development of highly scalable computational approaches capable of testing spatial associations among hundreds of thousands to millions of gene pairs. Second, several of the tools introduced in this manuscript rely on strong underlying assumptions and are therefore applicable only to cell-based spatial transcriptomic data. Specifically, they are based on the following key hypotheses:

1. The distribution of transcripts across a homogeneous cell population can be modeled by a negative binomial or Poisson distribution.
2. RNA concentration, rather than transcript count, is the primary determinant of cell state; consequently, RNA concentration is assumed to be homogeneous across similar cell types.

These assumptions limit the applicability of our gene selection and data normalization methods to datasets generated by protein-based multiplexed imaging technologies, such as CODEX, cyclic immunofluorescence, or imaging mass cytometry [Moffitt et al.]. As these technologies continue to gain traction, there is a pressing need to develop new gene (or protein) selection and normalization strategies specifically tailored to protein imaging data, grounded in robust and validated statistical frameworks. It is worth highlighting that more than a decade ago, foundational work was been done to characterize the theoretical distribution of protein abundance across cells [Karmakar and Bose; Friedman et al.; Cohen et al.]. While integrating these theoretical insights with the inherently complex and noisy data generated by multiplexed imaging remains a significant challenge, doing so holds great promise. We believe that bridging this gap could substantially enhance our ability to analyze protein imaging data and, ultimately, to unravel the intricate spatial organization of biological tissues.

## Data and code availability

All datasets used in this manuscript were downloaded from websites and publicly available repositories (see Methods). TranspaceR is available on the following GitHub repository: https://github.com/BOSTLAB/TranspaceR.

## Acknowledgements

We thank the members of the Bost lab for their suggestions and comments, as well as Alice Balfourier, Leanne De Koning and Nicolas Servant for their highly constructive feedbacks on the manuscript and Pierre Gestraud and Marc Gabriel for their help on the R package.

## Author contributions

F.M. wrote the pipeline, performed the analysis and developed the methodologies. P.B. developed the methodologies, performed the analysis and wrote the manuscript.

## Methods

In this manuscript, analysis were performed using R 4.3.1, Python 3.10.16 (for Scimilarity analysis) and 3.9.0 (for zarr file extraction).

### Datasets used in the manuscript

Throughout this manuscript we used the following already published datasets :

- The Xenium human lymph node dataset, downloaded from 10X Genomics website (https://www.10xgenomics.com/datasets/human-lymph-node-preview-data-xenium-human-multi-tissue-and-cancer-panel-1-standard).

- The CosMX human prefrontal cortex dataset, downloaded from Nanostring website (https://nanostring.com/products/cosmx-spatial-molecular-imager/ffpe-dataset/human-frontal-cortex-ffpe-dataset/).

- The CosMX human pancreas dataset, also downloaded from Nanostring website (https://nanostring.com/products/cosmx-spatial-molecular-imager/ffpe-dataset/cosmx-smi-human-pancreas-ffpe-dataset)

- The MERFISH human tonsil dataset, from [Zhao et al.]. The dataset was downloaded from the Gene Expression Omnibus repository (https://www.ncbi.nlm.nih.gov/geo/query/acc.cgi?acc=GSM8649279).

- The Xenium human liver dataset, downloaded from 10X Genomics website (https://www.10xgenomics.com/datasets/human-liver-data-xenium-human-multi-tissue-and-cancer-panel-1-standard).

- The CosMX human liver dataset, downloaded from Nanostring website (https://nanostring.com/products/cosmx-spatial-molecular-imager/ffpe-dataset/human-liver-rna-ffpe-dataset/).

### Reanalysis of the museum of spatial transcriptomic

We downloaded the latest updated version of the Museum of Spatial Transcriptomic database (as of December 2024) from the following link: https://docs.google.com/spreadsheets/d/1sJDb9B7AtYmfKv4-m8XR7uc3XXw_k4kGSout8cqZ8bY. With regard to the scope of this paper, we limited our analysis to the ‘smFISH’ tab.

### Data loading

To extract the count matrix and associated metadata from the Xenium object, we utilized the .zarr files. A Python script was developed, using the zarr, pandas, and scipy.sparse Python libraries. First, we accessed the ‘cells.zarr’ file to extract cell ID information and cell metadata. The count matrix, stored as a sparse matrix in the ‘cell_feature_matrix’ dataset, was reconstructed using the ‘csr_matrix’ function (scipy.sparse module). Then we retrieved corresponding gene names from the ‘transcripts.zarr’ file. The dataframes created for metadata and count matrix were saved as csv files. For CosMx datasets, a Seurat object containing the data was available. The gene count matrix and negative probes data are retrieved from their layers and combined into a single data frame. The obtained expression data is saved as a CSV file. From the Seurat object, the cell metadata layer is saved into its own CSV file.

### Low quality gene and cell filtering

Lowly expressed genes were filtered out by first computing a threshold using Otsu’s binarization method using an in-house implementation where for each of the 100 percentiles of the data vector, the weighted intravariance resulting from the binarization using the percentile value is computed, before selecting the percentile value with the lowest associated intravariance. Before applying Otsu’s threshold, the total gene count were first log transformed in order to obtain a data distribution that is more gaussian-like.

For cell filtering, a lower and upper limit for cell size is determined *a priori* based on the estimated cell composition of the studied tissue and the table of the minimal and maximal size for various cell types (Figure S1A, Supplementary Table 1). In practice we never used a lower limit below a radius of 2µm and an upper radius limit above 15µm. In addition to a size-based cell filtering, we also implemented a filtering based on the total amount of detected RNA molecules within the cell: we simply use the threshold of 1000 Unique Molecular Identifier (UMI) commonly used in the field of scRNA-seq and scaled it proportionally to the size of the gene panel relative to the total number of genes (set to 20.000).

### Reference cell size

When available, reference cell size data were obtained from the BioNumbers database (bionumbers) [Milo et al.]. For monocytes, endothelial cells and hepatocytes the data were respectively extracted from the following publications: [Schmid-Schönbein et al.], [Rubin et al.] and [Rohr et al.]. Cell radius was computed using the following relationship between a sphere radius **r** and **its** volume V:

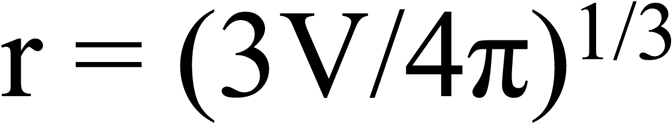

When available, the standard deviation of the cell volume was converted to the standard deviation of the cell radius using the error propagation formula:

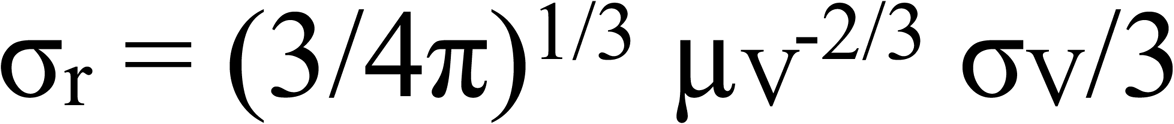

Where µ_V_ and σ_V_ are the mean and standard deviation of the cell volume.

### Statistically variable gene selection

For the detection of genes displaying an excess of variance, we assumed that gene expression values were following a negative binomial distribution which over-dispersion parameter was shared by all genes. The shared over-dispersion θ parameter was first estimated through a simple linear regression, implemented in the lm() R function from the **stats** package, thanks to the link between the mean µ and the variance σ^2^ of a negative binomial distribution:

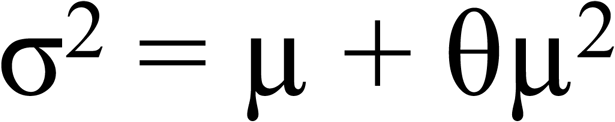

The variance score is then computed by taking for each gene the ratio between the observed variance and the theoretical variance computed using the estimated θ and the equation above. In order to estimate for each gene the variance of its variance estimator we used the following strategy:

- We first split the µ values into 31 bins by computing 30 corresponding quantiles µ_i_ using the quantile() R function.

- For each of the µ_i_ values, we sample N_cell_ times from a negative binomial distribution of parameter µ_i_ and θ using the rnbinom() function of the **MASS** package, compute the empirical variance and repeat N times in order to estimate the variance of the estimator. By default N is set to 500. N_cell_ corresponds to the total number of cells within the studied dataset.

- We then estimate the estimator variance for each gene through linear interpolation: we used the approx() function from the stats package and performed the interpolation after log transformation of the mean and of the estimator variance.

If the number of cells is too high (by default above 10.000 cells), an asymptotic estimation of estimator variance for each quantile value is performed by computing the estimator variance for a list of small N_cell_ values (by default 1000, 1500, 2000, 2500, 3000, 5000, 10000) before performing a linear model between the estimator variance and the log N_cell_. The final estimator variance corresponding to the real N_cell_ is then computed using the predict() R function.

Benefiting from the fact that according to the central limit theorem the variance estimator is following a gaussian distribution of which the mean and the standard deviation parameters are known, we can compute a p-value for each of the genes using the pnorm() function, before performing a Benjamini Hochberg multiple testing correction step using the p.adjust() base R function.

We use a similar approach for the detection of genes with an excess of zeros but instead we use the following property of a random variable X following a negative binomial distribution:

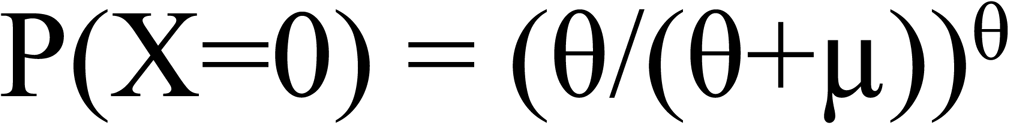

In that case the θ parameter is estimated using the above equation and the nls() R function. We used the nl2sol algorithm (algorithm=‘port’) and set the initial value through the moment’s method by taking the average observed value of (σ^2^ -µ)/µ^2^. The associated p-values are computed in a similar fashion to the variance estimator, with the only difference being that after each sampling of the negative binomial distribution we compute the proportion of counts equal to zero and not the variance. The excess zero score is defined as the difference between the observed proportion of zeros and the theoretical one.

In the manuscript we considered that a gene was significantly variable if its corrected p-value is below 0.01.

### Spatially variable gene selection

Spatially variable gene selection was performed using Geary’s C index. First we computed a Delaunay triangulation using the delaunay() function from the **RCDT** package before converting to an adjacency matrix using the graph_from_edgelist() and as_adjacency_matrix() functions from the **igraph** package. The adjacency matrix is then symmetrized and converted to a sparse matrix object (dgCMatrix object from the R Matrix package),. In order to efficiently compute Geary’s index we reformulated its computation as a matrix multiplication problem. Indeed if we note W the adjacency matrix and x the gene expression vector, the numerator of the Geary’s C index is equal to:

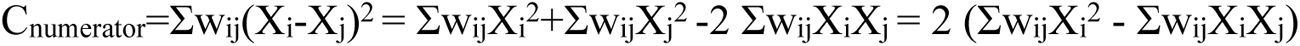

The numerator can thus be written as C_numerator_=2 (W.x^2^-x^T^.W.x) where x^T^ is the transpose of the gene expression vector and x^2^ the gene expression vector which entires have been raised to the square. As both the spatial neighborhood adjacency matrix and the gene expression vectors are highly sparse, the computation speed is thus significantly improved. The p-value is computed using the formula provided in the excellent book of Cliff and Ord [Cliff and Ord]. As we realized that this book is not easily accessible, we provide the formula for the Geary’s C estimator variance in the case where the expression vector does not follow a gaussian distribution:

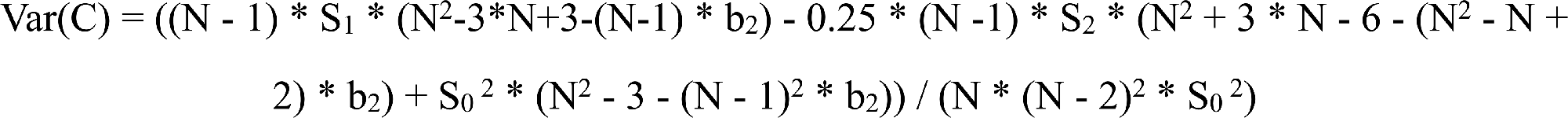

Where N is the number points/cells, S_0_ the sum of all spatial neighborhood matrix entries, S_1_ the sum of all squared entries of the neighborhood matrix, S_2_ the total sum of the row and column sums squared and b_2_ the kurtosis of the gene expression vector computed using the kurtosis() function from the **e1071** R package. The corresponding p-value was finally estimated by assuming a normal distribution of the estimator and using the known mean (E(C) = 1) and standard deviation of the estimator and computed using the pnorm() function. P-values were corrected for multiple testing by the Benjamini Hochberg method using the p.adjust() function

### Variogram computation by FFT

Our implementation is based on the work of Marcotte [Marcotte] to which have a added a step of rasterization. We first split the image tissue region into squares and sum the gene expression of all cells within the square. If no cell is present within a square, the expression value is set to NA. While the size of the squares can be set manually, by default it is computed by taking the mean of the tissue width and length and divide it by 200.

Once the initial 2D expression matrix **f** of size n_x_ x n_y_ has been computed for each gene we can compute their corresponding ‘variogram map’ **Γ**. We note I the indicator matrix, i.e. a matrix of size n_x_ x n_y_, which entries are set to 1 if cells were present in the corresponding squares and to 0 if not (i.e. if f entry is NA). Similarly we note **f^2^** the matrix of size n_x_ x n_y_ which entries are the squared entries of f. Finally we note TF() and TF**^-1^**() the Fourrier transform and inverse Fourrier transform. As described in [Marcotte] we have:

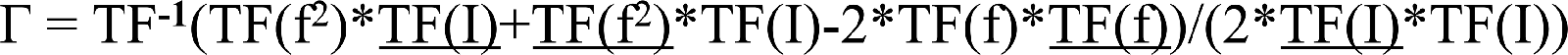

Where * represents element wise matrix multiplication (i.e. Hadamard product) and is the complex conjugate.. The Fourrier transforms are computed using the fft() function with default parameters with inverse transforms are computed using the fft() function with the inverse parameter set to TRUE.

The last part consists in converting this variogram map into a regular one dimensional variogram: for each entry/pixel of Γ, we compute its euclidean distance to the central pixel before finally computing the mean entry values of pixels with the same distance.

Once the empirical variogram has been computed, we fit for each gene three different variogram models :

- A constant model such that γ(r) = N

- An exponential model such that γ(r) = C (1-exp(-r/τ)) + N

- An fine-tuned exponential model such that γ(r) = C (1-exp(-(r/τ)^α^)) + N

Each model is fitted using the nls() function with initial parameter values being set to 0 and the algorithm parameter to ‘port’. In order to compare the different models, we computed their respective AIC score, assuming a gaussian error/likelihood. The model with the lowest AIC is then selected. A gene was considered as being spatially variable if its associated model was not the constant one, that more than 50% of the variance can be explained spatially, i.e. that C/(C+N) > 0.5, and that the theoretical variogram has an associated R^2^ bigger than 0.85.

### Differential gene expression analysis and simulation

Non-parametric differential gene expression was performed using the kruskal.test() function from the R **stats** package. The glm-Poisson regression was performed using the glm() R function with the parameter family set to “poisson” and other parameters set to default, thus corresponding to a Poisson glm with a logarithmic link function. For each test, two models were built: one including only an intercept and one including both an intercept and a group effect. The two models were compared using a likelihood ratio test implemented in the anova() function with the parameter test set to ‘Chisq’. Data simulation was performed using the rgamma() function for the sampling of volume values, with the shape parameter set to 6.4 and the rate to 0.12, in order to have a similar volume distribution as for the cells from the CosMx human pancreas dataset.

### Data simulation for dimensionality reduction benchmarking

In order to simulate a realistic dataset made of 10 distinct cell clusters of 1000 cells each, we used the following approach:

- For each cell we simulated a random volume value drawn from a gamma distribution of shape parameter set to 6.5 and shape parameter set to 0.1.

- For each cell cluster, we sample a random number of marker genes from a Poisson distribution of mean parameter set to 10.

- For each of the marker gene we considered that its expression within its associated cell cluster followed a Poisson distribution which mean parameter is the product of the cell volume and a given concentration parameter. This concentration parameter is randomly sampled from a list of values extracted from the Xenium human lymph node sample by fitting a Poisson mixture model to each gene. This is done using the flexmix() function from the **flexmix** package with default parameters.

### Dimensionality reduction and clustering

SVD was performed using the irlba() function from the **irlba** package while NMF was performed using the nmf() function of the **RcppML** R package. In both cases default parameters were used except the number of latent dimension which was set to 20 in the case of the Xenium human lymph node sample and 10 for the simulation data. In addition before performing SVD or NMF, each gene variance was set to 1 by dividing the expression vector by its standard deviation. The glm-pca was performed using the init_glmpca_pois() and fit_glmpca_pois() functions of the **fastglmpca** package. For the init_glmpca_pois() function, the parameters col_size_factor and row_intercept were set to TRUE and the number of components to 20 for the Xenium human lymph node sample and 10 for the simulation data, while for the fit_glmpca_pois() function the maximal number of iterations was set to 10. For the Poisson NMF we used the fit_poisson_nmf() function from the **fastTopics** package with the number of component also set to 20 for the Xenium human lymph node sample and 10 for the simulation data. Both the glm-pca and Poisson NMF methods were applied to the raw unscaled data.

### Metrics for clustering methods comparison

Normalized Mutualized Information (NMI) was computed using the NMI() function from the **aricode** package with default parameters. Adjusted Rand Index (ARI) was computed using the ARI() function from the **aricode** package with default parameters. The PanglaoDB cell marker database [Franzén et al.] was downloaded from the PanglaoDB server (www.panglaodb.se). Gene sets which contain less than 5 genes expressed in the lymph node dataset were removed before per forming any analysis. Gene Set Enrichment Analysis [Subramanian et al.] was performed using the iterative.bulk.gsea() function from the **liger** R package.

For the computation of the incompatible expression score we first computed the mean expression of each measured gene in each cell cluster. We then scale each gene mean expression value by dividing it by the maximal value across cell clusters.

### Automated cell annotation

Deep-learning based cell annotation is performed using Scimilarity (version 0.3.0). We developed a command-line Python script that accepts a CSV expression matrix as input and generates two output files. The script utilizes the **scanpy** Python library to read and preprocess the query expression matrix using the read_csv() function. A pre-trained Scimilarity model was loaded via the CellAnnotation() function from the **Scimilarity** module. The query dataset was aligned with the model using the align_dataset() function from scimilarity.utils. The aligned data was then normalized using the lognorm_counts() function from the same module. Cell type predictions are generated by applying the get_predictions_knn() function to the aligned dataset. This function returns predicted labels, indices of the nearest neighbors, distances to the nearest neighbors, and statistical metrics of the prediction. The predicted cell types and their corresponding statistical metrics were then saved to a CSV file. We compute the counts of each predicted cell type and save this information to a separate output file. To assess the confidence of the annotations, we calculate the mean selected annotation index relative to all other indices. A pandas dataframe was finally constructed to store the predictions and their statistical metrics. The mean confidence annotation indices for each cell type was also calculated.

For the annotation using CellTypist, the expression data were imported into Python using the **scanpy** library. The data were preprocessed to replace missing values with zeros using np.nan_to_num() function. Normalization to a total count of 10,000 was performed with sc.pp.normalize_total() . A logarithmic transformation is applied with the sc.pp.log1p() function. The pre-trained model Immune_All_Low.pkl was used ton annotate the Xenium human lymph node dataset and was retrieved at https://www.celltypist.org/models. For the CosMX human frontal cortex dataset, the Adult_Human_PrefrontalCortex.pkl model was used and retrieved similarly. The predictions were made using the celltypist.annotate() function, with majority voting option enabled. The annotated data is converted into an AnnData object and exported as a CSV file.

---

**Supplementary Figure 1:**
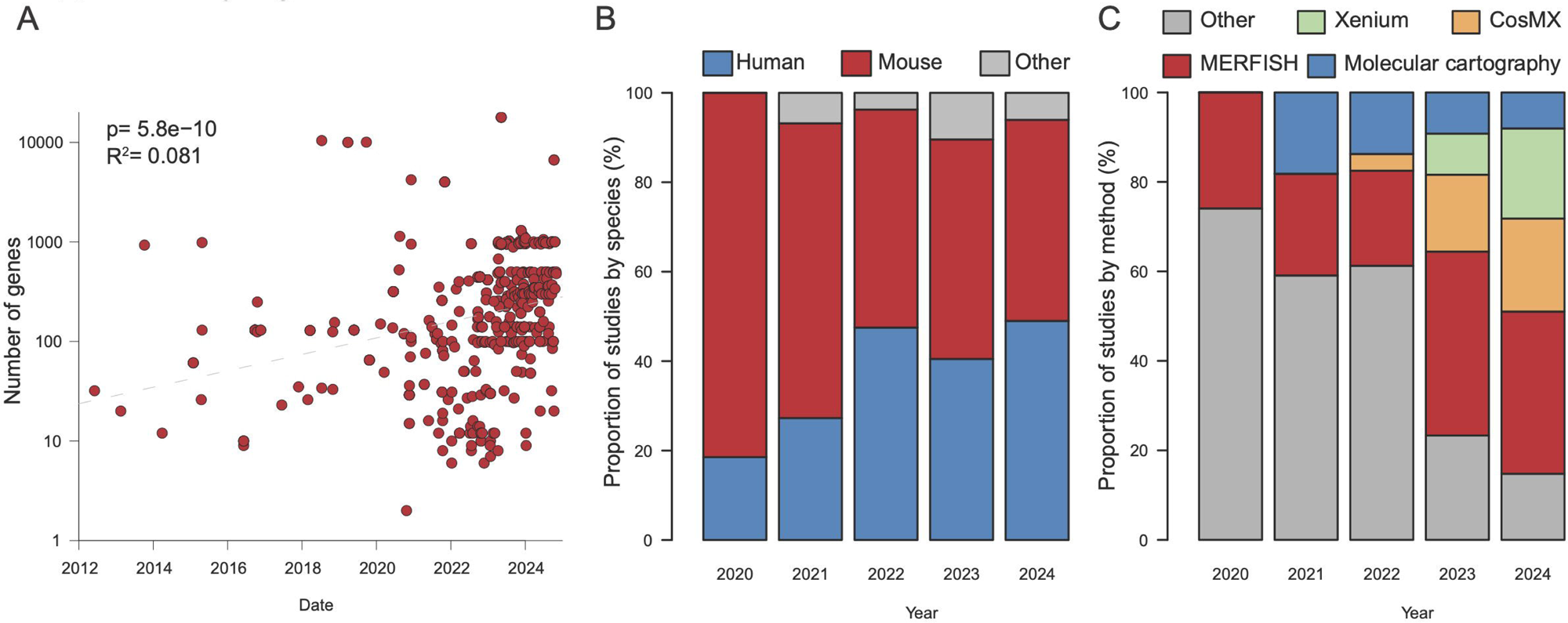
**(A)** Number of genes measured in multiplexed smFISH datasets according to publication year. The grey dashed line corresponds to a linear fit in the logarithmic space. Statistical significance was assessed using a Likelihood Ratio Test. **(B)** Proportion of multiplexed smFISH datasets containing samples from human, mouse or other species across time. **(C)** Proportion of multiplexed smFISH datasets generated using the Xenium, CosMX, MERFISH, Molecular cartography or other technologies across time.

**Supplementary Figure 2.**
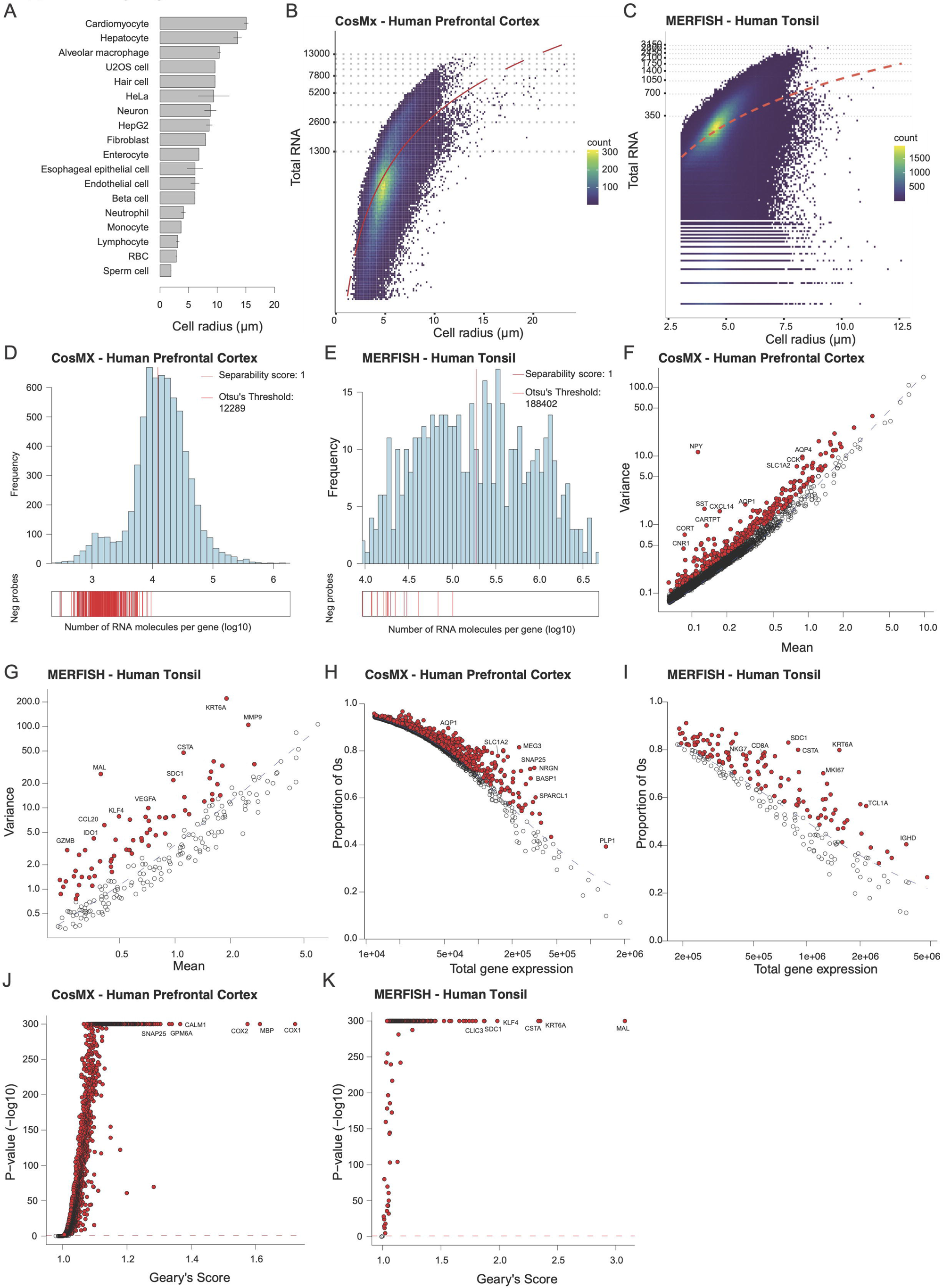
**(A)** Cell radius of various cell types. Black lines correspond to twice the standard deviation when available. **(B)** Relationship between cell radius and total RNA molecules detected in the CosMX human prefrontal cortex dataset. The red dashed line corresponds to a quadratic fit. **(B)** Relationship between cell radius and total RNA molecules detected in the CosMX human prefrontal cortex dataset. The red dashed line corresponds to a quadratic fit. **(C)** Relationship between cell radius and total RNA molecules detected in the MERFISH human tonsil dataset. The red dashed line corresponds to a quadratic fit. **(D)** Distribution of the number of RNA molecules detected for each gene in the CosMX human prefrontal cortex dataset. The vertical red line corresponds to the threshold computed using Otsu’s method. **(E)** Distribution of the number of RNA molecules detected for each gene in the MERFISH human tonsil dataset. The vertical red line corresponds to the threshold computed using Otsu’s method. **(F)** Relationship between gene expression mean and variance in the CosMX human prefrontal cortex dataset. The dashed blue line corresponds to the theoretical relation between gene expression mean and variance according to the fitted negative binomial model. Genes displaying a significant excess of variance (corrected p-value <0.01) are colored in red. **(G)** Relationship between gene expression mean and variance in the MERFISH human tonsil dataset. The dashed blue line corresponds to the theoretical relation between gene expression mean and variance according to the fitted negative binomial model. Genes displaying a significant excess of variance (corrected p-value <0.01) are colored in red. **(H)** Relationship between gene expression mean and proportion of zeros in the CosMX human prefrontal cortex dataset. The dashed blue line corresponds to the theoretical relation between gene expression mean and proportion of zeros according to the fitted negative binomial model. Genes displaying a significant excess of zeros (corrected p-value <0.01) are colored in red. **(I)** Relationship between gene expression mean and proportion of zeros in the MERFISH human tonsil dataset. The dashed blue line corresponds to the theoretical relation between gene expression mean and proportion of zeros according to the fitted negative binomial model. Genes displaying a significant excess of zeros (corrected p-value <0.01) are colored in red. **(J)** Geary’s C score for gene expression in the CosMX human prefrontal cortex dataset. Genes displaying a significantly high Geary’s C score (corrected p-value <0.01) are colored in red. (**K**) Geary’s C score for gene expression in the MERFISH human tonsil dataset. Genes displaying a significantly high Geary’s C score (corrected p-value <0.01) are colored in red.

**Supplementary Figure 3.**
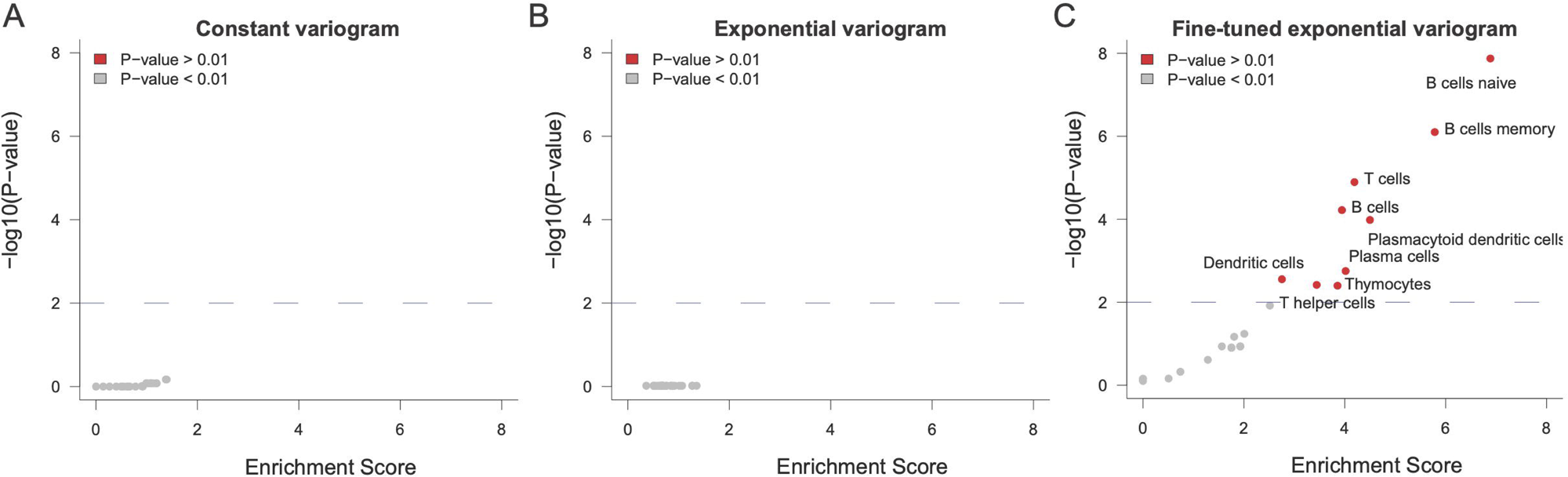
**(A)** Gene set enrichment analysis of the genes categorized as having a constant variogram in the Xenium human lymph node dataset. Genes displaying a significantly enrichment (corrected p-value <0.01) are colored in red. **(B)** Gene set enrichment analysis of the genes categorized as having an exponential variogram in the Xenium human lymph node dataset. Genes displaying a significantly enrichment (corrected p-value <0.01) are colored in red. (**C)** Gene set enrichment analysis of the genes categorized as having a fine-tuned exponential variogram in the Xenium human lymph node dataset. Genes displaying a significantly enrichment (corrected p-value <0.01) are colored in red.

**Supplementary Figure 4:**
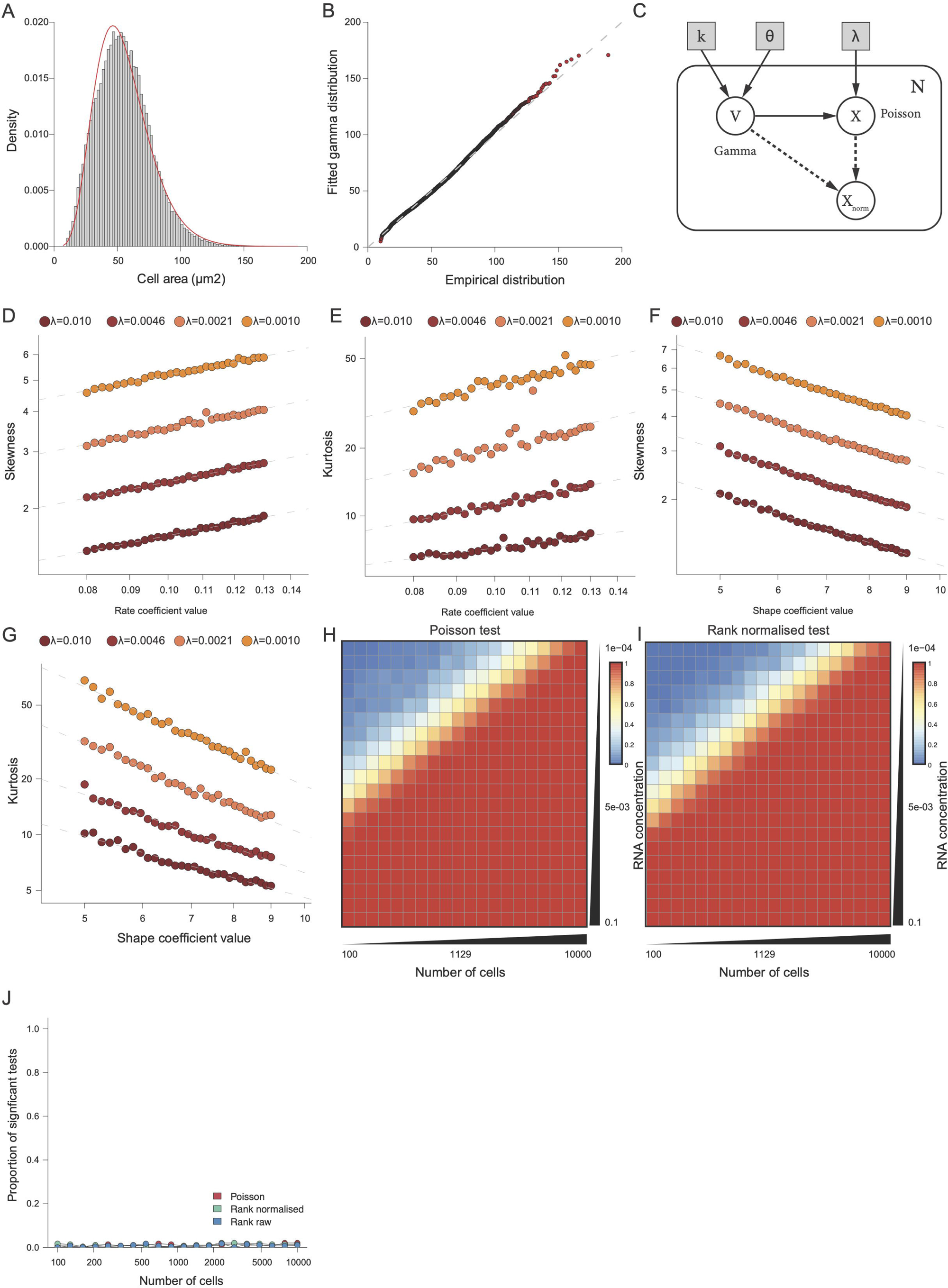
**(A)** Distribution of cell size in the CosMX human pancreas dataset. The red curve corresponds to the fitted gamma distribution. **(B)** Q-Q plot comparing of the empirical and fitted gamma distribution of cell area. **(C)** Plate model of the simulation model. **(D)** Relationship between the skewness of synthetic volume-normalized data as a function of the rate coefficient value. Each dashed grey line corresponds a linear regression for a different RNA concentration. **(E)** Relationship between the kurtosis of synthetic volume-normalized data as a function of the rate coefficient value. Each dashed grey line corresponds a linear regression for a different RNA concentration. **(F)** Relationship between the skewness of synthetic volume-normalized data as a function of the shape coefficient value. Each dashed grey line corresponds a linear regression for a different RNA concentration. **(G)** Relationship between the kurtorsis of synthetic volume-normalized data as a function of the shape coefficient value. Each dashed grey line corresponds a linear regression for a different RNA concentration. **(H)** Impact of cell number and RNA concentration on the statistical power of Poisson regression. **(I)** Impact of cell number and RNA concentration on the statistical power of rank test performed on volume normalized data. **(J)** False positive analysis of Poisson regression and of rank test on raw and volume normalized data

**Supplementary Figure 5:**
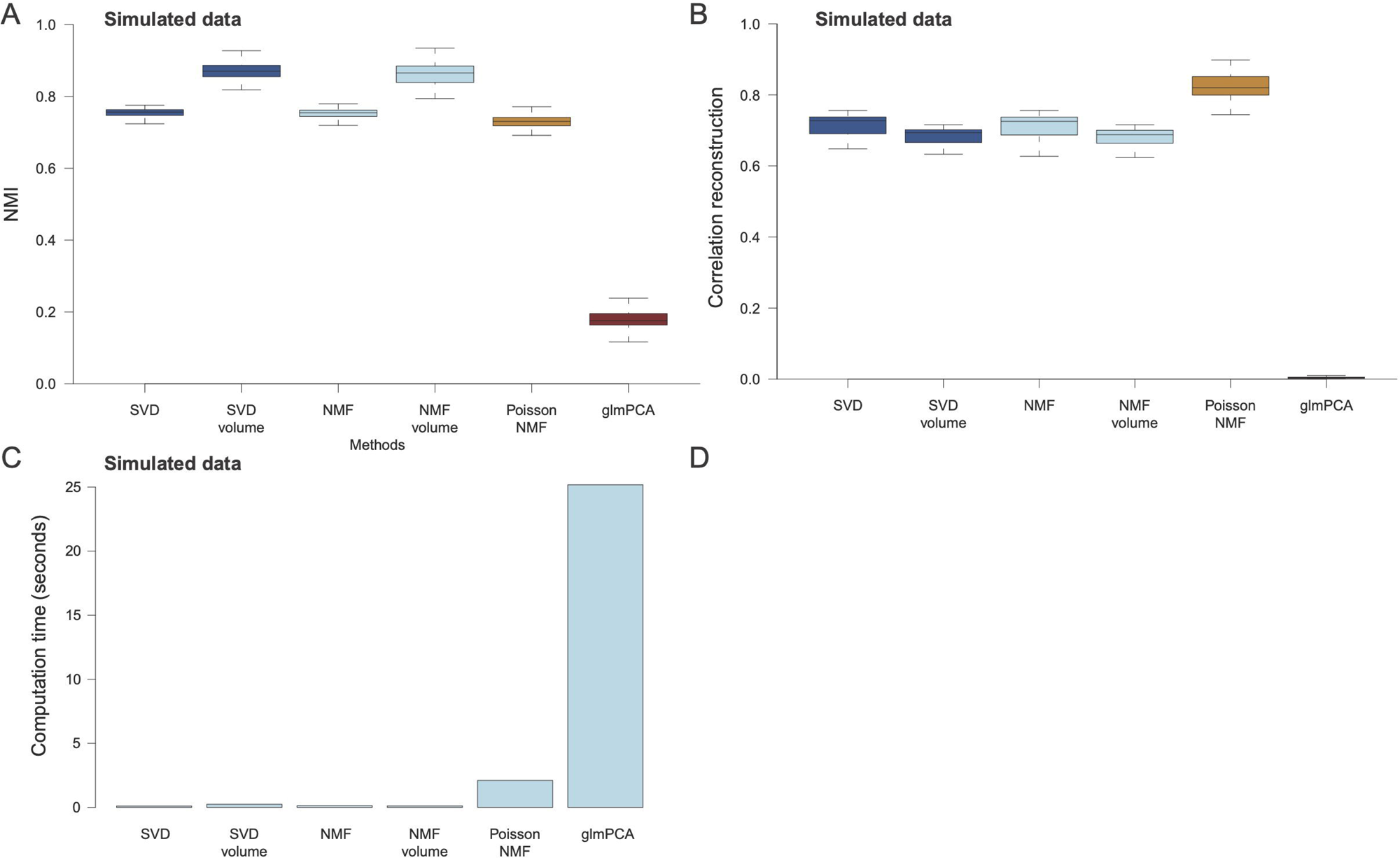
**(A)** Distribution of the Normalized Mutual Information (NMI) between inferred and reference clusters according to the dimensionality reduction technique used when applied to simulated data. The thick line corresponds to the median, and the bottom and upper limits of the box correspond to the first and third quartiles, respectively. The lower and upper whiskers correspond to the lowest and highest values, respectively, within the range of the first and third quartiles ±1.5 times the IQR. The boxplot is based on N=300 simulations. **(B)** Distribution of the correlation between the original and reconstructed data according to the dimensionality reduction technique used when applied to simulated data. The thick line corresponds to the median, and the bottom and upper limits of the box correspond to the first and third quartiles, respectively. The lower and upper whiskers correspond to the lowest and highest values, respectively, within the range of the first and third quartiles ±1.5 times the IQR. The boxplot is based on N=300 simulations. **(C)** Average computation time according to the dimensionality reduction technique used when applied to simulated data.

**Supplementary Figure 6:**
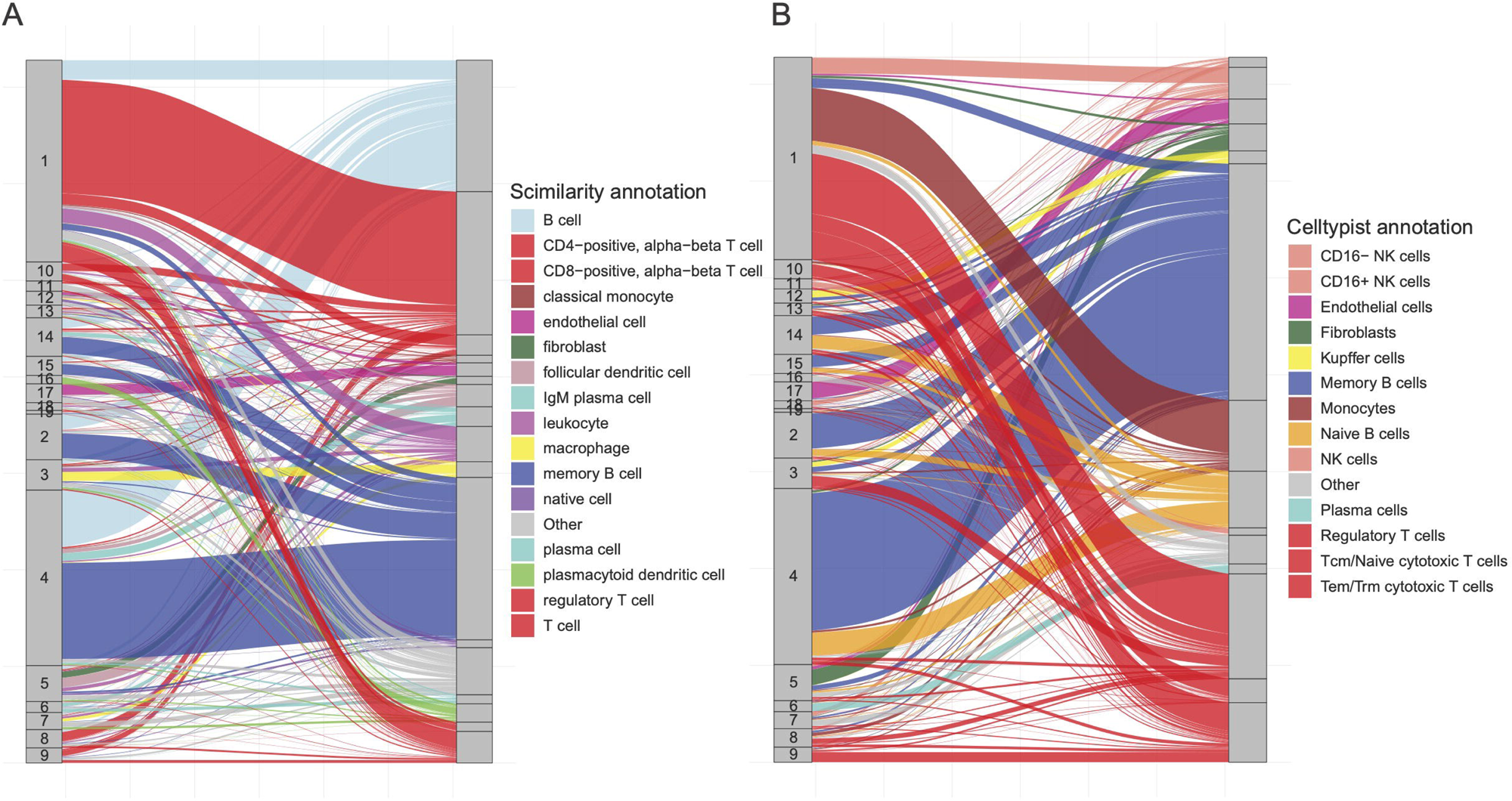
**(A)** Comparison of Scimilarity annotation and unsupervised clustering. **(B)** Comparison of CellTypist annotation and unsupervised clustering.

