## Supplementary Note 1 for "Next generation statistical framework for next generation spatial transcriptomics data"

### Normalisation of smFISH data

April 14, 2025

#### 1 Analysis of the variance source

Let $X$ be the number of RNA molecules of a given gene observed in a given cell. There are strong empirical evidences that the number of RNA molecules is proportional to the volume of the cell volume, i.e. that the RNA concentration is constant in a homogeneous cell population Padovan-Merhar et al. (2015). Then if the concentration of the RNA $\lambda$ and the cell volume $v$ are known we can further assume that the distribution of RNA transcripts detected in a cell follows a Poisson distribution of parameter $\lambda v$:

$$X|\lambda, v \sim \mathcal{P}oisson(\lambda v) \tag{1}$$

As the volume of a cell is strictly positive and following empirical observations, we further consider that $v$ follows a Gamma distribution:

$$v \sim \mathcal{G}amma(k, \theta) \tag{2}$$

Where $k$ and $\theta$ are two positive parameters corresponding to the shape and scale parameters of the distribution. The marginal distribution of $X$ is thus a negative binomial distribution, sometimes called the Gamma-Poisson distribution.

By the law of total variance we have:

$$Var(X) = \mathbb{E}(Var(X|\lambda v)) + Var(\mathbb{E}(X|\lambda v)) \tag{3}$$

the left and right parts of the equation respectively corresponding to the unexplained and explained part of the variance by the latent paremeter $\lambda v$. Using the properties of the Poisson distribution we obtain :

$$\begin{aligned} Var(X) &= \mathbb{E}(\lambda v) + Var(\lambda v) \\ &= \lambda \mathbb{E}(v) + \lambda^2 Var(v) \\ &= \lambda k\theta + \lambda^2 k\theta^2 \end{aligned} \tag{4}$$

So the ratio of variance explained $V_{explained}$ by the cell size is equal to :

$$V_{explained}(\lambda, \theta) = \frac{\lambda\theta}{1 + \lambda\theta} \tag{5}$$

Therefore the correlation between the cell volume and the RNA count is simply :

$$r_{volume}(\lambda, \theta) = \sqrt{\frac{\lambda\theta}{1 + \lambda\theta}} \tag{6}$$

### 2 Interpretation

The final equation allows us to understand the global behavior of gene expression data obtained by smFISH technologies:

- If the RNA concentration is high or the cell volume varies widely, then the main source of variation is the variation of the cell volume.
- In the other case, the majority of gene expression variation simply comes from Poisson sampling of the transcripts.

Importantly, the RNA concentration parameter $\lambda$ considered here is not the actual RNA concentration but rather the observed one that can be seen as the real RNA concentration multiplied by the detection efficiency of the protocol used. For instance, highly multiplexed smFISH technologies, such as the commercial Xenium platform, have a low detection rate due to the use of complex *in-situ* amplification steps like Rolling Circle Amplification (RCA) Janesick et al. (2022).
